# Computational analysis of lamin isoform interactions with nuclear pore complexes

**DOI:** 10.1101/2020.04.03.022798

**Authors:** Mark Kittisopikul, Takeshi Shimi, Meltem Tatli, Joseph R. Tran, Yixian Zheng, Ohad Medalia, Khuloud Jaqaman, Stephen A. Adam, Robert D. Goldman

**Affiliations:** Department of Cell and Developmental Biology, Feinberg School of Medicine, Northwestern University, Chicago, IL; Department of Biophysics, UT Southwestern Medical Center, Dallas, TX; Cell Biology Center and World Research Hub Initiative, Institute of Innovative Research, Tokyo Institute of Technology, Yokohama, Japan; Department of Biochemistry, University of Zurich, Zurich, Switzerland; Department of Embryology, Carnegie Institution for Science, Baltimore, MD; Lyda Hill Department of Bioinformatics, UT Southwestern Medical Center, Dallas, TX

## Abstract

Nuclear lamin isoforms form fibrous meshworks associated with nuclear pore complexes (NPCs). Using data sets prepared from sub-pixel and segmentation analyses of 3D-Structured Illumination Microscopy images of WT and lamin isoform knockout mouse embryo fibroblasts, we determined with high precision the spatial association of NPCs with specific lamin isoform fibers. These relationships are retained in the enlarged lamin meshworks of *Lmna*^*-/-*^ and *Lmnb1*^*-/-*^ fibroblast nuclei. Cryo-ET observations reveal that the lamin filaments composing the fibers contact the nucleoplasmic ring of NPCs. Knockdown of the ring-associated nucleoporin ELYS induces NPC clusters that exclude lamin A/C fibers, but include LB1 and LB2 fibers. Knockdown of the nucleoporins TPR or NUP153 alter the arrangement of lamin fibers and NPCs. Evidence that the number of NPCs is regulated by specific lamin isoforms is presented. Overall the results demonstrate that lamin isoforms and nucleoporins act together to maintain the normal organization of lamin meshworks and NPCs within the nuclear envelope.

## Introduction

The nuclear envelope (NE) is a complex multicomponent structure separating the nuclear genome from the cytoplasm. It has evolved as a highly compartmentalized multifunctional organelle with a wide range of functions. The NE structure includes the nuclear lamina (NL), a double membrane bilayer forming a lumen continuous with the endoplasmic reticulum and nuclear pore complexes (NPCs). However, details of the structural and spatial relationships among the components of the NE have been difficult to define. This lack of information is largely attributable to the dense packing and close spatial relationships of the structures comprising the NE (Aebi et al., 1986; Fisher et al., 1986; Goldman et al., 1986; McKeon et al., 1986). To better understand the structural relationships within the NE, we have combined super-resolution light microscopy with recently developed computer vision techniques. This approach has allowed us to quantitatively analyze the structural organization of the lamins and NPCs in the NE by making highly precise measurements of lamin structures and NPC localization over large areas of the NE. Our goal is to test the utility of large data sets to provide new insights into the interactions between these two major components of the NE.

The four major lamin isoforms in somatic cells are lamin A (LA), lamin C (LC), lamin B1 (LB1), and lamin B2 (LB2). These type V intermediate filament proteins are closely apposed to the inner nuclear membrane where they assemble into discrete fibrous meshworks. In mouse embryo fibroblast nuclei, the NL is a 13.5 nm thick layer composed of 3.5 nm diameter filaments (Turgay et al., 2017). Using three-dimensional structured illumination microscopy (3D-SIM) combined with computer vision analysis, we demonstrated that bundles of these filaments, termed fibers in the light microscope, are non-randomly organized into complex interwoven meshworks within the NL (Shimi et al., 2015; Turgay et al., 2017). Notably, each lamin isoform assembles into distinct meshworks with similar structural organization (Shimi et al., 2015). Previous studies on *Lmna*^*-/-*^ MEFs (Sullivan et al., 1999) showed that loss of lamin A/C caused dramatic changes in nuclear morphology with some NPC clustering. Subsequently, we showed that the meshworks formed by individual lamin isoform fibers are significantly expanded in size in *Lmna* or *Lmnb1* knockout (KO) MEF nuclei compared to the lamin meshworks in WT or *Lmnb2* KO MEF nuclei demonstrating that LA/C and LB1 interactions are required for normal lamin fiber meshwork structure in WT MEFs (Shimi et al., 2015).

The NPCs penetrate the NE forming transport passageways delineated by the fusion of the inner and outer nuclear membranes, thereby allowing for bidirectional transport across the NE. They are composed of multiple copies of 30 proteins known as nucleoporins (Beck and Hurt, 2016). For many years, it has been apparent that there are structural interactions between the NL and NPCs of vertebrate nuclei. The earliest studies on identification of components of the NE identified a cell free NPC-NL fraction that could be isolated under fairly stringent conditions suggesting their strong physical association (Kay et al., 1972; Dwyer and Blobel, 1976; Scheer et al., 1976; Aebi et al., 1986). In addition, both lamins and the NPCs are relatively immobile in the plane of the NE indicating that both are anchored in some fashion (Broers et al., 1999; Moir et al., 2000; Rabut et al., 2004). Both the nuclear lamins and NPC structures are closely associated with chromatin at the nuclear periphery (Guelen et al., 2008; Peric-Hupkes et al., 2010; Ibarra and Hetzer, 2015) with the NPCs located in spaces where both the lamina and heterochromatin appear to be discontinuous (Fawcett, 1966; Ou et al., 2017). Super-resolution microscopy analysis of lamins and NPCs in *Lmna*^*-/-*^ fibroblasts also found NPCs closely associated with exogenously expressed LA and LC in Xie et al. (2016). Some clustering of NPCs within the remaining LB1 networks has also been reported in *Lmna*^*-/-*^ fibroblasts (Xie et al., 2016). Our previous study by cryo-ET also supports the close association of lamin filaments with the NPCs (Turgay et al., 2017; Tatli and Medalia, 2018).

Biochemical analyses of lamin-NPC interactions have shown connections between lamins and a subset of specific nucleoporins (Hase and Cordes, 2003; Krull et al., 2004; Al-Haboubi et al., 2011).More recently, proximity-dependent biotin identification, BioID, recognized several lamin-associated nucleoporins including Nup153, ELYS and TPR (Roux et al., 2012; Xie et al., 2016). These nucleoporins localize to the nucleoplasmic aspect of NPCs which lie in close proximity to the NL (Walther, 2001; Rasala et al., 2008). The distribution of NPCs is nonrandom with characteristic center to center spacing varying according to species ranging from human to frog (Maul, 1977). Furthermore, removal of all lamins from mouse MEFs or mESC derived fibroblast-like cells leads to clustering of the NPCs, which can be rescued by re-expression of either A or B-type lamins (Guo and Zheng, 2015). These observations suggest that lamins play an important role in regulating the distribution of NPCs.

Although the extant evidence strongly suggests that lamins interact with nucleoporins to anchor the NPCs in the NE, how each lamin isoform contributes to these interactions remains unknown. In this study, we investigate the structural relationships between each lamin isoform fiber meshwork and NPCs over large areas of the NE at nanoscale precision using 3D-SIM with newly developed computational procedures for sub-pixel quantitative image analysis. The analysis involves collecting positional information derived from large numbers of individual NPCs and determining their spatial relationship to each lamin isoform fiber comprising the NL meshworks. This quantitative approach is necessitated by the complexity of the four lamin fiber meshworks and NPCs located within a thin layer at the nuclear surface. The results of our analyses demonstrate that NPCs are closely associated with lamin fibers. At higher resolution cryo-ET confirms that both LA/C and LB1 filaments interact closely with the NPCs at the nucleoplasmic ring. Targeted disruption of nucleoporins and lamin isoforms demonstrates the interdependence of the spatial distributions of lamin fibers and NPCs.

## Results

### NPCs are structurally linked to lamin fibers

We used 3D-SIM and image reconstruction to determine the structural relationships among immunolabeled lamin fiber meshworks and NPCs in MEFs. NPCs in WT MEFs were distributed all across the NL region, but did not show an obvious co-localization with any of the lamin meshworks, as indicated by the very few white areas in merged overlays (Figure 1A). This was remarkable because some co-localization of lamins and NPCs would be expected by chance given the densely packed environment of the NL. This lack of co-localization between lamins and NPCs suggested the existence of a bona fide spatial relationship. We took advantage of our previous finding that the spaces or “faces” delineated by lamin fibers comprising the meshworks increase in size in *Lmna*^*-/-*^ and *Lmnb1*^*-/-*^ MEF nuclei (Shimi et al., 2015). This allowed us to examine the association between NPCs and specific lamin isoforms in WT, *Lmna*^*-/-*^, and *Lmnb1*^*-/-*^ MEFs. Importantly, NPCs remained in close proximity to the LA and LB1 fibers in the expanded meshworks of *Lmna*^*-/-*^ and *Lmnb1*^*-/-*^ MEF nuclei and were absent in the meshwork faces (Figure 1B). These results strongly suggest that LA and LB1 are required for the normal distribution of NPCs. Although these images provide qualitative evidence that there is an association between lamin isoform fibers and NPCs, it is important to verify such associations using a quantitative approach to ascertain the extent of the relationships between each lamin isoform fiber and NPCs.

**Figure 1.**
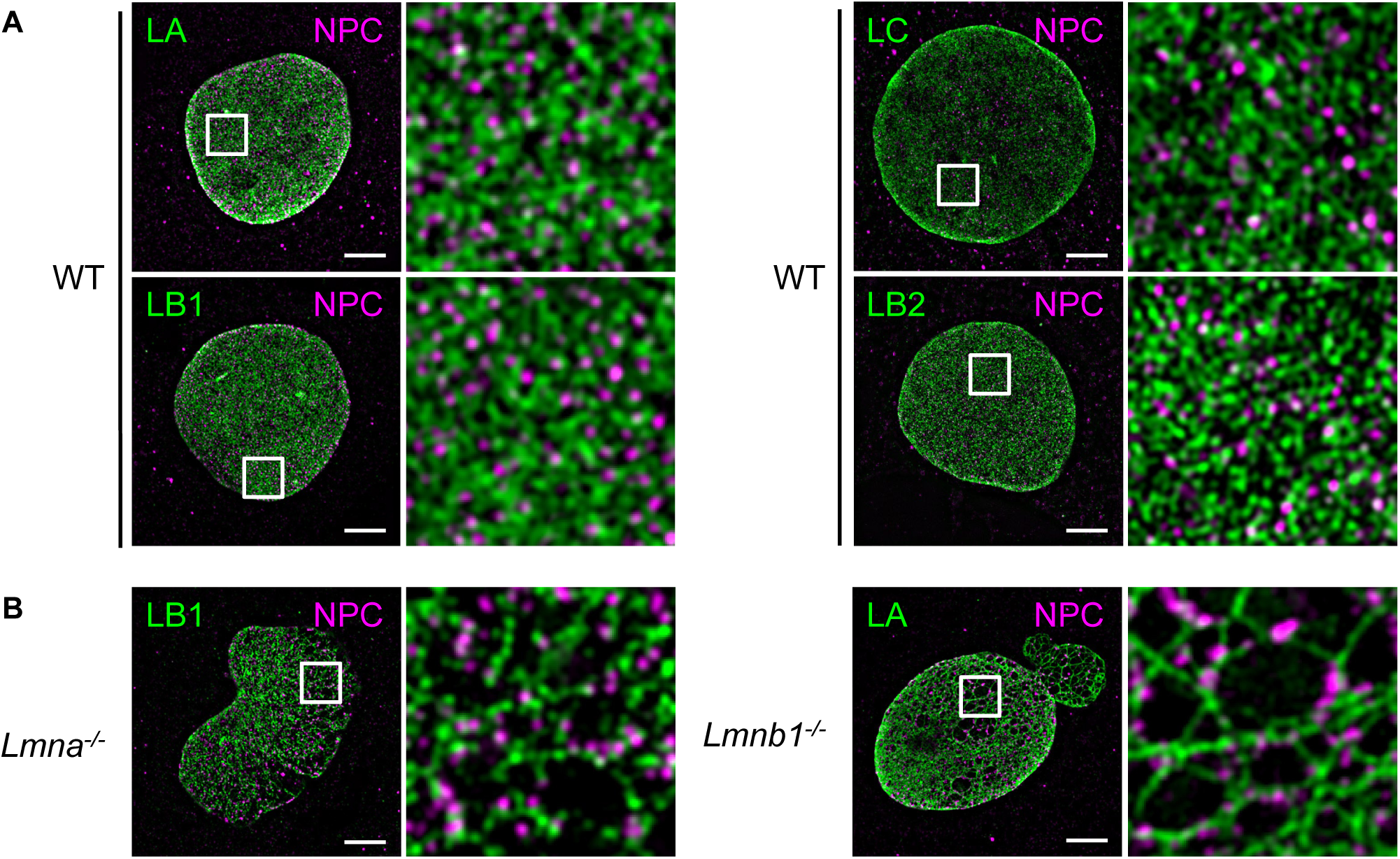
NPCs are arranged along LA and LB1 fibers in enlarged lamin meshworks. Colabeling of lamins and nuclear pore complexes in WT and lamin KO MEF nuclei using indirect immunofluorescence with a pair of specific antibodies against each lamin isoform (LA, LB1, LB2 or LC) and the FXFG-repeat nucleoporins. A) WT MEF nuclei colabeled with the indicated lamin isoform and FXFG-repeated nucleoporins. B) Nuclei of *Lmna*^*-/-*^ (left pair) and *Lmnb1*^*-/-*^ (right pair) MEFs. The indicated areas with white squares are enlarged approximately eight-fold along each edge and displayed on the right side of each pair of images. Scale bar = 5 *µm*.

### Image analysis reveals enrichment of NPCs within 30 to 100 nm of LA fibers in WT and *Lmnb1*^*-/-*^ MEFs

We developed quantitative image analysis tools to precisely determine the spatial relationships between lamin isoform fibers and NPCs, and to localize both structures with sub-pixel precision in dense and sparse lamin meshworks (Figure 2A; details of analysis tools in Materials and Methods). We reasoned that by measuring the distances between the centers of lamin fibers and the center of lamin meshwork faces to the centers of NPCs (Figure S1), we could quantitatively assess the association of NPCs with individual lamin isoforms. To evaluate the frequency of observing distances between the lamin fibers or face centers and NPCs by chance, we compared our observed distance measurements to the expected distances under a null hypothesis, which assumes the NPCs and lamin meshworks have no relationship and are thus independently distributed. For example, we measured the LA fiber center to NPC center distance in WT cells as compared to the expected distances assuming no relationship (Figure 2B compare the measured data in the blue violin plot on top vs the expected distances in the red violin plots on bottom). By examining the difference in the observed from the expected distributions (Figure 2C), we could see a paucity (green) or excess (purple) of NPCs at certain distances from the centers of LA fibers. For example, in a single WT nucleus we observed fewer NPCs within 30 nm of the fibers and an excess of NPCs between 30 and 100 nm relative to the null hypothesis (green area; Figure 2C WT). In order to validate this approach, we performed the same analysis of the LA fiber to NPC distance in a single *Lmnb1*^*-/-*^ MEF nucleus (Figure 2B). As in the WT nucleus, we saw an excess of NPCs between 30 and 100 nm in the *Lmnb1*^*-/-*^ nucleus (Figure 2C). This agreed with the qualitative observation that the NPCs were associated with, but not co-localized with lamin fibers (Figure 1A, B, 2A).

**Figure 2.**
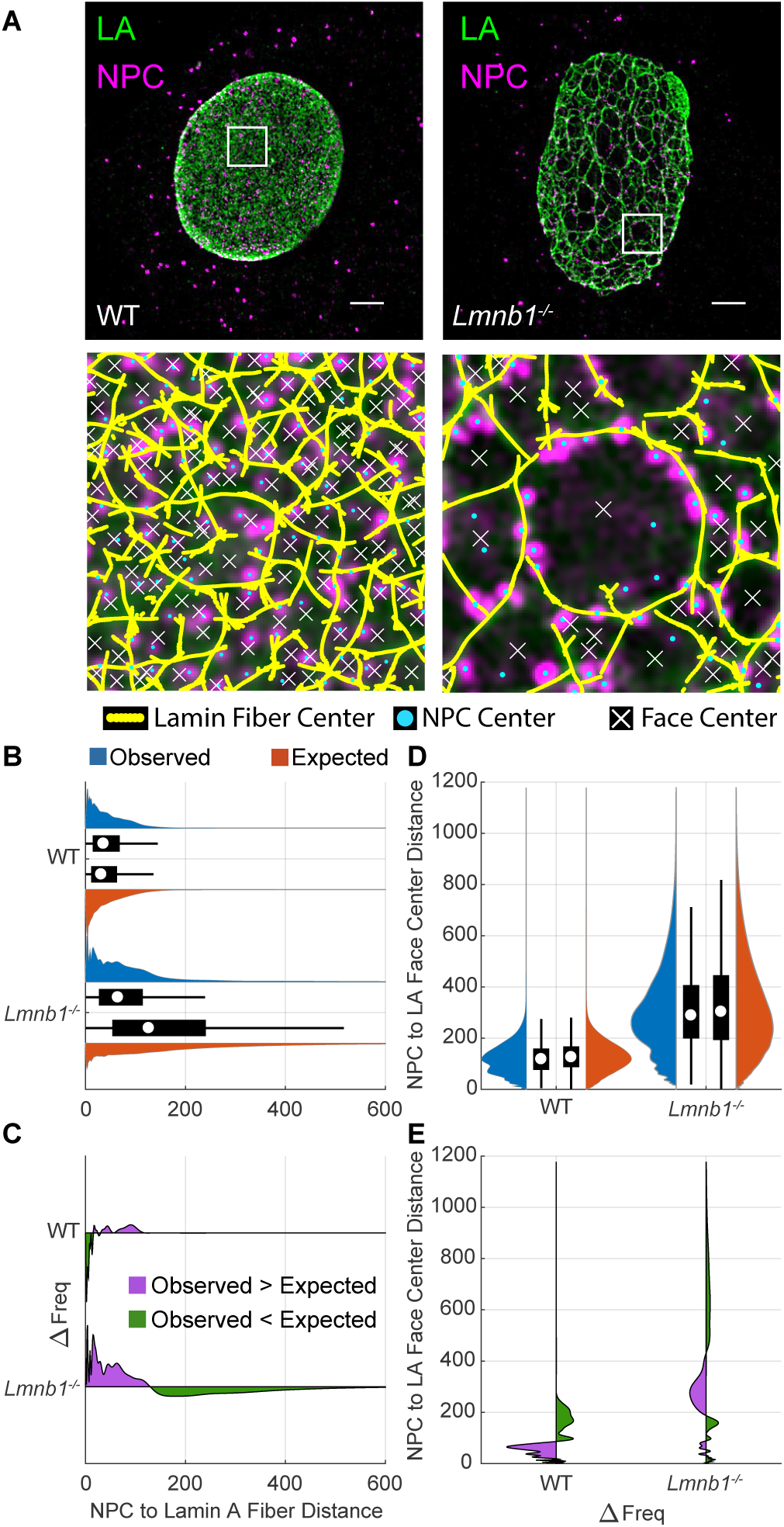
Computational image analysis reveals that NPCS are closely associated with LA fibers. A) Immunofluoresence images labeling LA (green) and NPCs (magenta) of WT and *Lmnb1*^*-/-*^ MEF nuclei as in Figure 1 were subjected to computational image analysis. White boxes in the top row are magnified 8 times along each edge. The centers of LA fibers (yellow dots), NPCs (cyan dots), and faces (white Xs) were segmented to subpixel precision (Kittisopikul et al. (2020); Appendix 1). Scale bar is 5 *µm*. B) Paired violin and box plots of NPC to LA fiber distances for the nuclei in (A). The violin (blue) and box plots on top represent the observed distance distributions. The violin (red) and box plots on bottom represent the expected distance distributions under the null hypothesis. The white circle indicates the median. The thick black bar indicates the interquartile range (IQR). The black whiskers indicate 1.5 times the IQR. C) Frequency difference plot of observed minus expected LA fiber to NPC distances. The green portion below the line indicates where the observed frequency is less than expected. The purple portion above the line indicates where the observed frequency is greater than expected. D) NPC to LA face center distances displayed as in (B), rotated 90 degrees counterclockwise. E) Frequency difference plot of NPC to LA face center distances, displayed as in (C), rotated 90 degrees counterclockwise.

Measuring the distance from the lamin face centers to NPCs allowed us to more precisely determine how NPCs are related to the lamin fibers. The faces are delineated by the lamin fibers composing the lamin isoform meshwork (Figure 2A; Shimi et al. (2015)). Their centers are points that are locally the most distant from the lamin fibers. This analysis also allowed us to account for changes in face size such as the enlargement seen in *Lmnb1*^*-/-*^ or *Lmna*^*-/-*^ nuclei (Figures 1B, 2A). Measuring both the distances of the NPCs to the lamin fibers and the centers of the faces, allowed us to examine a 2D bivariate statistical distribution in a single nucleus (Figure S1). To explore if the NPCs also had a relationship with the center of the faces, we found the points the most distant from the lamin fibers within a local area (white Xs, Figure 2A). For a circle, this would be the center, but other shapes may have multiple centers (see Methods). We measured the distances between the center of the NPCs and the center(s) of the faces (Figure S1G) and then compared that distribution to the null hypothesis (Figure 2D, E). In both the WT and the *Lmnb1*^*-/-*^ nucleus, we observed median distances that were smaller than expected. This means that the NPCs were closer to the center of the faces than expected by chance. This is consistent with the observation that NPCs did not directly colocalize with the lamin fibers, but had a lateral proximal relationship.

We combined the distances of the NPCs to the lamin fibers and the distances of the NPCs from the face centers into two-dimensional histograms to represent the bivariate distribution (Figure S2). The two-dimensional histograms showed that there was an expectation that NPCs would be near the LA fibers and away from the faces by chance in a broad distribution. However, the NPCs were offset from the LA fibers in a narrower than expected distribution (Figure S1A-F). In the WT MEFs, the negative correlation between the distances was also apparent, which is expected since the NPCs that are farther from the lamin fibers tend to be closer to the face centers (Figure S1A-B). However, the two-dimensional histograms of single nuclei were sparse and noisy indicating that additional distance measurements were needed for evaluation.

The localizations of both lamin fibers and NPCs were based on finding local maxima within the continuous reconstruction of the fluorescence intensity from critically sampled 3D-SIM images and was not dependent on rounding to the nearest pixel (See Methods and Supplement; Kittisopikul et al. (2020)). Here we focused on localizing lamin fibers and NPCs resolved by 3D-SIM, and not their specific molecular components consisting of individual 3.5 nm lamin filaments Turgay et al. (2017) and/or specific nucleoporins. Furthermore, we measured the distance between structures localized within two channels separated by their chromatic properties, and thus these distance measurements were not limited by resolution (Stelzer, 1998). The main limitations to the precision of the localization and distance measurements are the inaccuracy of indirect immunofluoresence labeling, signal-to-noise ratio, and structured illumination microscopy reconstruction artifacts. This was mitigated by examining the distribution of tens of thousands of distance measurements. These analyses permitted us to express the magnitude of differences in the co-distributions, or the lack thereof, in terms of nanometers with high statistical power (see Appendix 1).

### LA and LB1 fibers have a more pronounced relationship with NPCs than LC and LB2 fibers in WT MEFs

We previously found that the four main lamin isoforms (LA, LC, LB1, and LB2) form independent meshworks (Shimi et al., 2015), and we sought to see if each isoform had a distinct relationship with NPCs. Having established our approach to analyzing lamin-NPC associations, we measured the distances between the center of individual NPCs and the center of the nearest lamin fiber across the surface of the nucleus closest to the coverslip of 10 WT nuclei for each lamin isoform. Overall, the data obtained supports the lack of direct colocalization between NPCs and lamin fibers, which we observed qualitatively and quantitatively in single nuclei (Figures 1, 2). The median distances from the centers of NPCs to the centers of LA fibers (40.4 nm; p < 0.001; Table 1A, Figure 3A, Figure S2A) and to the centers of LB1 fibers (38.1 nm; p < 0.001; Table 1A, Figure 3A) were similar. The observed median distances were 6 nm greater than the expected distribution (+6.9 nm LA; +6.0 nm LB1; Table 1A, Figure 3A, B; Figure S2C). The expected distribution represents the distances between NPCs and lamins that we would expect under the null hypothesis that there is no relationship between the position of NPCs and lamins. It was calculated by a Monte Carlo simulation randomly placing a NPC within the segmented area of the nucleus. The median distance between NPCs and the center of faces in the LA meshworks was similar (119.3 nm; −11.7 nm vs expected; p < 0.001; Table 1B) to LB1 (118.3 nm; −10.8 nm vs expected; p < 0.001; Table 1B) and both median distances were less than expected if the lamins and NPCs were not associated (Figure 3C; Table 1B). These data show that NPCs and LA or LB1 fibers are not directly colocalized, but have a proximal lateral relationship. These findings suggest that NPCs and LA or LB1 fibers are structurally linked within the NL.

**Table 1A:** Lamin fiber - NPC center to center distance distributions for WT, Lmnb1-/-, and Lmna-/- MEFs.

| Cell Genotype | Lamin Labeled | Observed (nm) |  | Expected (nm) |  | Obs. - Exp. (nm) |  | P-Values |  | Num. of NPCs N |
| --- | --- | --- | --- | --- | --- | --- | --- | --- | --- | --- |
|  |  | Median | St. Dev. | Median | St. Dev. | Median | St. Dev. | Median | St. Dev. |  |
| wt | LA | 40.4 | 38.0 | 33.5 | 56.5 | 6.9 | -18.5 | 0.00 |  | 14780 |
| wt | LC | 32.8 | 35.0 | 32.1 | 49.4 | 0.7 | -14.5 | 0.37 | 0.01 | 11459 |
| wt | LB1 | 38.1 | 36.2 | 32.1 | 56.9 | 6.0 | -20.7 | 0.00 |  | 15150 |
| wt | LB2 | 27.6 | 29.2 | 28.1 | 38.7 | -0.6 | -9.6 | 0.00 |  | 17146 |
| Lmnb1-/- | LA | 45.1 | 48.6 | 42.4 | 216.8 | 2.7 | -168.2 | 0.59 | 0.00 | 11971 |
| Lmna-/- | LB1 | 34.9 | 34.5 | 35.8 | 297.7 | -0.8 | -263.1 | 0.00 |  | 9740 |

Median and standard deviation of the observed and expected lamin fiber to NPC center to center distances, the difference between them, p-values (see Methods), and number of NPCs. The data in each row was collected from 10 cells. The Mann Whitney U test and Ansari-Bradley test were used as in described in the Materials and Methods. P-values in red were above the Bonferroni corrected alpha value of $0.05/8 \text{ tests} = 0.006$ .

**Table 1B:** Face - NPC center to center distance distributions for WT, Lmnb1-/-, and Lmna-/- MEFs.

| Cell Genotype | Lamin Labeled | Observed (nm) |  | Expected (nm) |  | Obs. - Exp. (nm) |  | P-Values |  | Num. of NPCs N |
| --- | --- | --- | --- | --- | --- | --- | --- | --- | --- | --- |
|  |  | Median | St. Dev. | Median | St. Dev. | Median | St. Dev. | Median | St. Dev. |  |
| wt | LA | 119.3 | 62.6 | 130.9 | 78.3 | -11.7 | -15.7 | 0.00 |  | 14780 |
| wt | LC | 122.4 | 57.1 | 125.7 | 69.0 | -3.3 | -11.9 | 0.00 |  | 11459 |
| wt | LB1 | 118.3 | 56.8 | 129.1 | 76.0 | -10.8 | -19.2 | 0.00 |  | 15150 |
| wt | LB2 | 116.7 | 51.5 | 117.3 | 58.9 | -0.6 | -7.3 | 0.25 | 0.08 | 17146 |
| Lmnb1-/- | LA | 124.0 | 90.0 | 146.0 | 235.2 | -22.0 | -145.2 | 0.00 |  | 11971 |
| Lmna-/- | LB1 | 122.1 | 55.7 | 133.2 | 304.3 | -11.1 | -248.6 | 0.00 |  | 9740 |

Median and standard deviation of the observed and expected lamin face to NPC distances, the difference between them, p-values (see Methods), and number of NPCs. The data in each row was collected from 10 cells. The Mann Whitney U test and Ansari-Bradley test were used as in described in the Materials and Methods. P-values in red were above the Bonferroni corrected alpha value of $0.05/7 \text{ tests} = 0.007$ .

**Figure 3.**
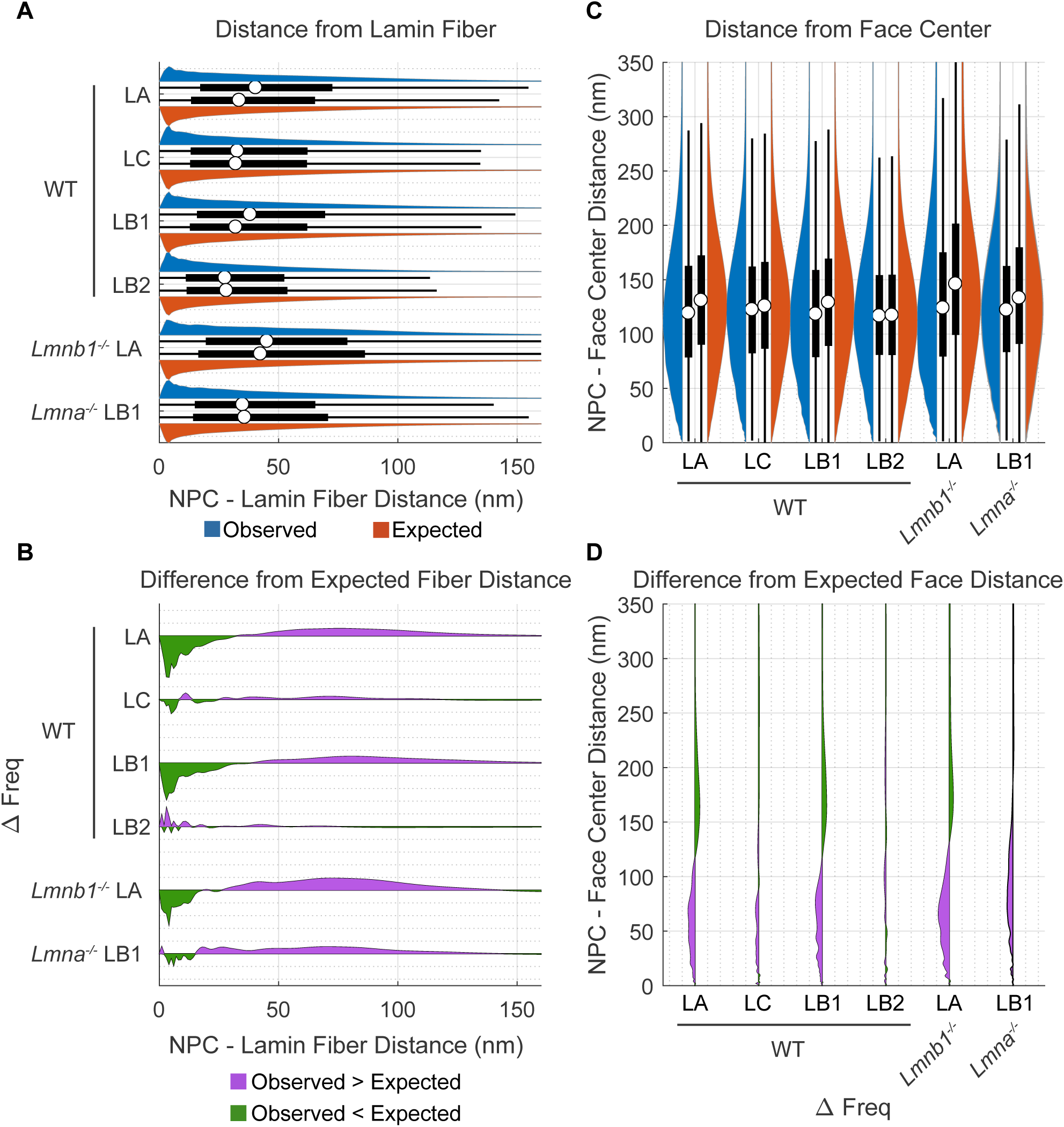
Quantitative analysis of Lamin-NPC distances over many nuclei reveals NPCs are offset from the center of LA and LB1 fibers in WT, *Lmna*^*-/-*^, and *Lmnb1*^*-/-*^ MEFs by 20 - 30 nm. A) Paired violin and box plots of NPC to lamin fiber distances. The violin (blue) and box plots on top represent the observed distance distributions. The violin (red) and box plots on bottom represent the expected distance distributions under the null hypothesis. The white circles indicate the medians. The thick black bar indicates the interquartile range (IQR). The black whiskers indicate 1.5 times the IQR. B) Frequency difference plots of observed minus expected lamin fiber to NPC distances. The green portion below the line indicates where the observed frequency is less than expected. The purple portion above the line indicates where the observed frequency is greater than expected. C) NPC to lamin face center distances displayed as in (A), rotated 90 degrees counterclockwise. D) Frequency difference plot of NPC to lamin face center distances, displayed as in (C), rotated 90 degrees counterclockwise.

In contrast to the relationships between the NPCs and LA or LB1, the median distance from LC fibers to NPC centers did not differ significantly from expected (32.8 nm observed, + 0.7 nm vs expected; p= 0.37; Table 1A, Figure 3A, Figure S2B). Also, the standard deviation of distances between LC fibers and NPCs (35.0 nm observed, −14.5 nm vs expected; p=0.01; Table 1A, Figure 3A) was not significant when using a Bonferonni corrected alpha level. While the p-value of 0.01 is smaller than the traditional alpha level of 0.05, we conducted multiple comparisons and thus need to compensate for Type I error. The Bonferroni correction of the alpha level across the 12 pairs of distributions compared in Tables 1A and 1B leads to an alpha level of 0.05/12 ≈ 0.004. However, the median distance determined for the NPC center to LC face center differed from the expected distribution (122.4 nm observed, −3.3 nm vs expected; p < 0.001; Table 1B, Figure 3C). While these measurements followed a pattern similar to that detected for LA and LB1, the magnitude of the differences were much smaller for LC (Figure 3C, D, Table 1B). Overall, these data suggested that the offset between NPCs and LC fibers is closer (median: 32.8 nm) than between NPCs and LA or LB1 fibers (medians: 40 nm). However, given the small differences in the LC fiber to NPC center measurements relative to expected, we cannot completely reject the null hypothesis for the LC fiber to NPC distances.

The relationship between LB2 fibers and NPCs in WT MEFs differed from the other lamin isoforms. We observed a statistically significant difference in medians from expected distributions between the centers of LB2 fibers and NPCs (27.6 nm observed; −0.6 nm vs expected; p < 0.001; Table 1A, Figure 3A, Figure S2D). However, the shift was an order of magnitude less and in the opposite direction than observed for LA and LB1 fibers. The median distance from NPCs to LB2 face centers (116.7 nm observed; −0.6 nm vs expected; Table 1A, Figure 3C) was not significantly different from expected. These findings suggest that there is no obvious relationship between the distribution of LB2 fibers and NPCs, or if there is, it cannot be discerned in our analyses.

### Knocking out *Lmna* affects the LB1-NPC relationship more than knocking out *Lmnb1* affects the LA-NPC relationship

The results presented in the previous section showed a clear spatial relationship between both LA and LB1 fibers and NPCs in the dense meshworks of WT MEF nuclei. The removal of either LA/C or LB1 by gene knockout in MEFs leads to dramatic changes in the remaining lamin meshwork characteristics, most notably an increase in the lamin mesh size (Figure 1B and Shimi et al 2015). Because the lamin fibers have close structural relationships with NPCs, we next wanted to determine if these relationships are altered when the lamin meshwork structure changes.

We analyzed the spatial relationships between LA fibers and NPCs in 10 *Lmnb1*^*-/-*^ nuclei using the same quantitative methods applied to our studies of WT nuclei. In *Lmnb1*^*-/-*^ nuclei, there was a greater median distance between LA fiber centers and NPC centers than expected (45.1 nm observed; +2.7 nm vs expected; Table 1A, Figure 3A, Figure S3A), however, this shift in medians was not statistically significant (p = 0.59, Table 1A). Interestingly a statistical test comparing the standard deviations showed that the distributions are significantly different (48.6 nm observed; −168.2 nm vs expected; p < 0.001; Table 1A, Figure 3A, B). This reflects the long tail of the expected distributions, since under the null hypothesis some NPCs may appear in the middle of the faces of the enlarged LA meshworks, that is, farther away from the lamin fibers. The median distance of NPCs from the LA face centers was less than expected by a large magnitude (124.0 nm; −22.0 nm vs expected; p < 0.001; Table 1B; Figure 3C, D). This difference is due to the distribution of the offsets of the NPCs from the lamin fibers, which is larger than the expected offset distributions where more NPCs were closer to the lamin fibers. The observed distance distributions of WT and *Lmnb1*^*-/-*^ MEFs (Figure 3A) both differ from the expected distributions under the null hypothesis in a similar manner (Figure 3B). This indicates that, in *Lmnb1*^*-/-*^ nuclei, the proximal lateral relationship between LA fibers and NPCs remains although the median distance between LA fibers and NPCs increased by 5 nm. Overall, this suggests that the distance between the centers of LA fibers and NPCs does not depend strongly on the presence of LB1 fibers.

The results showed a relationship similar to LA fibers in WT MEFs for distances less than 30 nm where NPCs occurred less frequently than expected (green area; Figure 3B) and more frequently than expected around 50-100 nm (purple area; Figure 3B). This differed from the analysis of the single nucleus which consisted mostly of enlarged faces (Figure 2A), whereas most nuclei typically had a mix of small and large faces (Figure 1B).

Interestingly, the median distances between the centers of LB1 fibers and NPCs in *Lmna*^*-/-*^ MEFs matched the expected distribution (34.9 nm observed; −0.8 vs expected; p < 0.001; Table 1A, Figure 3 A,B, Figure S3B). Recall that in contrast, the LB1 fiber to NPC median distances in WT MEFs were slightly larger and differed from the expected (38.1 nm; p < 0.001; Table 1A, Figure 3A). Additionally, the difference between the frequencies of the observed and expected distributions were smaller in magnitude in *Lmna*^*-/-*^ MEFs compared to WT MEFs along with a small positive peak suggesting some colocalization (Figure 3B). The standard deviation of LB1 fiber to NPC medians in *Lmna*^*-/-*^ MEFs did differ significantly from expected (34.9 nm observed; −263.1 nm vs expected; p < 0.01; Table 1A, Figure 3A, B) reflecting the enlarged faces in *Lmna*^*-/-*^ MEFs. LB1 face center to NPC center distances were significantly different from expected with a large change in magnitude (122.1 nm observed; −11.1 nm vs expected; p < 0.001; Table 1B, Figure 3C, D). As in WT MEFs, this reflects a lateral proximal relationship between LB1 fibers and NPCs in *Lmna*^*-/-*^ MEFs.

The average number of NPCs per nucleus in a single focal plane closest to the coverslip was reduced to 1000 NPCs in *Lmna*^*-/-*^ MEFs compared to 1200 in *Lmnb1*^*-/-*^ MEFs and 1500 in WT MEFs (Table 1, Figure S4), suggesting that both LA and LB1 are involved in regulating NPC number.

### Cryo-electron tomography (Cryo-ET) and immunogold labeling show LA/C and LB1 filaments contacting the nucleoplasmic ring of NPCs

In order to further investigate the relationship between lamin filaments and NPCs, we carried out cryo-ET of WT MEFs coupled with immunogold labeling of both LA and LB1. We hypothesized that this may shed additional insights on the lamin-NPC interaction and could reflect the relative abundance of LA and LB1 filaments contacting the NPC. We extracted 340 nm x 340 nm x 20 nm subtomograms around the nucleoplasmic ring of NPCs (Figure 4A; Turgay et al. (2017)) and counted the number of LA/C or LB1 filaments (Figure 4B). We observed more LA/C filaments than LB1 filaments in these regions (Figure 4C). These results also demonstrate that both LA and LB1 fibers are closely associated with the nucleoplasmic ring.

**Figure 4.**
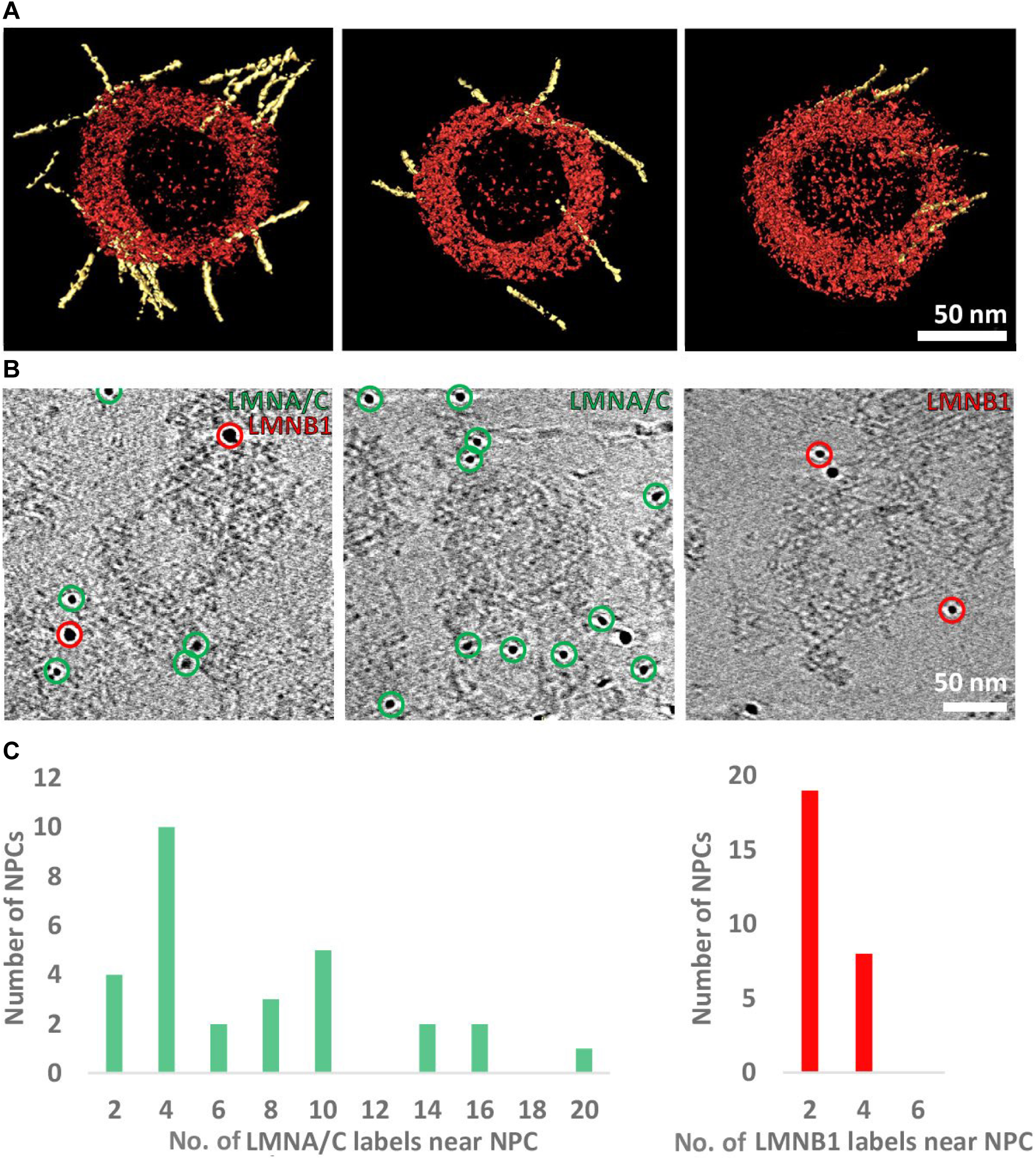
Cryo-electron tomography showing LA/C and LB1 filament contacts with the nucleoplasmic ring. A) Lamin filaments (yellow) interact with NPCs (red) as seen by surface rendering representations of cryo-sub-tomograms. B) Gold labelling of lamin filaments observed by cryo-ET. The position of Lamin A/C labels (green) and Lamin B1 labels (red) are indicated. Double labeling (left) or labeling of individual lamin isoforms were analyzed and presented as histograms. The unmarked gold particles (B-middle, right) are fiducial markers. C) A total number of 214 Lamin A/C labels and 70 Lamin B1 labels were detected around 47 nucleoplasmic rings.

### Organizational changes in LA meshworks and NPCs differ in response to silencing the expression of ELYS, TPR and NUP153

The cryo-ET observations, taken together with the demonstration that there was a proximal lateral association between NPCs and both LA and LB1 fibers suggested that there are attachments of lamin filaments to nucleoplasmic components of NPCs. We next explored the potential roles of individual nucleoporins in attaching lamin fibers to the NPCs. For these studies we focused on ELYS, NUP153 and TPR, all components of the nucleoplasmic NPC structures that are in close proximity to the lamina (Roux et al., 2012). The nucleoporin ELYS is a component of the nucleoplasmic ring of NPCs and is required for post-mitotic NPC assembly where it binds to the chromosomes and recruits the Nup107-160 complex of the nucleoplasmic ring (Franz et al., 2007). TPR and Nup153 are both components of the nuclear basket structure of the NPC tht associates with the nucleoplasmic ring (Duheron et al., 2014; Krull et al., 2004). We employed siRNA knockdown of each nucleoporin to determine their potential roles in linking the NPC to lamin fibers (Figure S6). We evaluated the efficacy of the knockdown by Western blot of whole cell lysates resulting in reductions of amount of each protein by 75%, 50%, or 40% for NUP153, ELYS, or TPR, respectively (Figure S5). Knockdown of either ELYS or TPR led to significant changes in NPC distribution and structural relationship to the LA fibers. The most dramatic effect was the reorganization of NPCs into clusters after ELYS knockdown (Figure 5A). Individual fluorescent puncta could still be resolved within each cluster indicating that some NPC structure was likely retained. In contrast, siRNA knockdown of NUP153 or TPR did not cause NPC clustering in WT MEFs (Figure 5A). The median distance between the centers of NPCs and LA fibers in ELYS depleted cells (70.8 nm; +20 nm vs scrambled; p < 0.001; Table 2A, Figure 5A, B, Figure S6) increased compared to scrambled siRNA controls (50.9 nm; p < 0.001; Table 2A, Figure 5A, B, Figure S6). Additionally, the median distance between face centers of the LA fiber meshwork and the NPCs was reduced (89.7 nm; Table 2B; Figure 5C) compared to scrambled siRNA (106.2 nm; p < 0.001; Table 2B, Figure 5C, Figure S6). These data suggested that LA fibers were being excluded from the ELYS depleted NPC clusters such that these clusters became located in large faces within the LA meshwork. Interestingly, the size of faces contained within the LA meshwork also appeared to increase upon ELYS knockdown (Figure 5A, F). As a measure of lamin face size, we summed the NPC to fiber distances and the NPC to face center distances, since, for a perfectly circular face in the meshwork, this quantity would be the radius of the circle with respect to each NPC. The face radius of the LA fiber meshwork (169.7 nm; Table 2C) significantly increased versus the scrambled siRNA control (163.3 nm; p < 0.001; Table 2C) upon ELYS knockdown indicating that the LA meshwork expanded when ELYS was depleted.

**Table 2A:** Lamin fiber to NPC center to center distance distributions of WT MEFs with TPR, NUP153, and ELYS knockdown.

| siRNA Knockdown | Lamin Labeled | Observed (nm) |  | Expected (nm) |  | Obs. - Exp. (nm) |  | P vs Exp. |  | Obs. - Scram. | P vs Scram. | Num. of NPCs N |
| --- | --- | --- | --- | --- | --- | --- | --- | --- | --- | --- | --- | --- |
|  |  | Median | St. Dev. | Median | St. Dev. | Median | St. Dev. | Median | St. Dev. |  |  |  |
| Scrambled | LA | 50.9 | 39.5 | 33.6 | 40.4 | 17.3 | -0.9 | 0.00 |  |  |  | 39096 |
| TPR KD | LA | 59.0 | 39.5 | 31.9 | 36.9 | 27.1 | 2.6 | 0.00 |  | 8.2 | 0.00 | 40767 |
| NUP153 KD | LA | 50.1 | 38.6 | 31.1 | 35.7 | 19.0 | 2.8 | 0.00 |  | -0.8 | 0.00 | 36066 |
| ELYS KD | LA | 70.8 | 48.9 | 32.9 | 42.4 | 37.9 | 6.5 | 0.00 |  | 20.0 | 0.00 | 21521 |
| Scrambled | LC | 42.9 | 36.1 | 31.7 | 42.6 | 11.2 | -6.5 | 0.00 |  |  |  | 37760 |
| TPR KD | LC | 56.6 | 38.1 | 31.2 | 54.4 | 25.4 | -16.2 | 0.00 |  | 13.7 | 0.00 | 35489 |
| NUP153 KD | LC | 39.9 | 35.1 | 29.8 | 35.6 | 10.1 | -0.5 | 0.00 |  | -3.0 | 0.00 | 39988 |
| ELYS KD | LC | 63.1 | 46.7 | 32.8 | 44.2 | 30.3 | 2.6 | 0.00 |  | 20.2 | 0.00 | 27053 |
| Scrambled | LB1 | 51.6 | 42.4 | 35.4 | 51.8 | 16.2 | -9.4 | 0.00 |  |  |  | 37383 |
| TPR KD | LB1 | 52.1 | 38.4 | 31.3 | 49.0 | 20.8 | -10.6 | 0.00 |  | 0.5 | 0.00 | 40899 |
| NUP153 KD | LB1 | 46.9 | 41.3 | 35.2 | 40.6 | 11.7 | 0.7 | 0.00 |  | -4.7 | 0.00 | 31145 |
| ELYS KD | LB1 | 48.5 | 40.1 | 31.1 | 40.6 | 17.4 | -0.5 | 0.00 |  | -3.1 | 0.00 | 24981 |
| Scrambled | LB2 | 30.1 | 33.8 | 34.4 | 67.2 | -4.4 | -33.4 | 0.00 |  |  |  | 35444 |
| TPR KD | LB2 | 28.6 | 30.3 | 30.2 | 75.0 | -1.7 | -44.7 | 0.00 |  | -1.5 | 0.00 | 36974 |
| NUP153 KD | LB2 | 25.6 | 30.9 | 32.3 | 39.9 | -6.6 | -9.0 | 0.00 |  | -4.4 | 0.00 | 31628 |
| ELYS KD | LB2 | 34.2 | 33.8 | 31.2 | 40.2 | 3.0 | -6.3 | 0.00 |  | 4.1 | 0.00 | 25215 |

Median and standard deviation of the observed and expected lamin fiber to NPC center to center distances, the difference between them, p-values (see Methods), and number of NPCs. The distributions were also compared to scrambled siRNA control. The data in each row was collected from 20 cells. The Mann Whitney U test and Ansari-Bradley test were used as in described in the Materials and Methods.

**Table 2B:** Lamin face to NPC center to center distance distributions of WT MEFs with TPR, NUP153, and ELYS knockdown.

| siRNA Knockdown | Lamin Labeled | Observed (nm) |  | Expected (nm) |  | Obs. - Exp. (nm) |  | P vs Exp. |  | Obs. - Scram. | P vs Scram. | Num. of NPCs N |
| --- | --- | --- | --- | --- | --- | --- | --- | --- | --- | --- | --- | --- |
|  |  | Median | St. Dev. | Median | St. Dev. | Median | St. Dev. | Median | St. Dev. |  |  |  |
| Scrambled | LA | 106.2 | 60.6 | 132.0 | 63.6 | -25.8 | -3.0 | 0.00 |  |  |  | 39096 |
| TPR KD | LA | 90.0 | 58.0 | 127.1 | 60.0 | -37.1 | -2.0 | 0.00 |  | -16.2 | 0.00 | 40767 |
| NUP153 KD | LA | 99.7 | 57.0 | 126.2 | 58.0 | -26.6 | -1.1 | 0.00 |  | -6.5 | 0.00 | 36066 |
| ELYS KD | LA | 89.7 | 58.8 | 129.7 | 64.4 | -39.9 | -5.6 | 0.00 |  | -16.4 | 0.00 | 21521 |
| Scrambled | LC | 109.1 | 58.1 | 126.5 | 65.2 | -17.4 | -7.2 | 0.00 |  |  |  | 37760 |
| TPR KD | LC | 89.9 | 55.6 | 125.8 | 73.4 | -35.9 | -17.7 | 0.00 |  | -19.2 | 0.00 | 35489 |
| NUP153 KD | LC | 106.6 | 55.5 | 122.9 | 57.7 | -16.3 | -2.2 | 0.00 |  | -2.5 | 0.00 | 39988 |
| ELYS KD | LC | 96.1 | 59.3 | 129.9 | 65.9 | -33.7 | -6.6 | 0.00 |  | -13.0 | 0.00 | 27053 |
| Scrambled | LB1 | 114.0 | 63.4 | 138.6 | 73.4 | -24.6 | -9.9 | 0.00 |  |  |  | 37383 |
| TPR KD | LB1 | 96.7 | 56.9 | 126.6 | 68.4 | -30.0 | -11.4 | 0.00 |  | -17.3 | 0.00 | 40899 |
| NUP153 KD | LB1 | 118.8 | 63.7 | 135.8 | 65.7 | -17.0 | -2.0 | 0.00 |  | 4.8 | 0.00 | 31145 |
| ELYS KD | LB1 | 101.5 | 58.2 | 125.6 | 62.2 | -24.1 | -4.0 | 0.00 |  | -12.5 | 0.00 | 24981 |
| Scrambled | LB2 | 138.8 | 59.7 | 134.6 | 85.8 | 4.2 | -26.1 | 0.00 |  |  |  | 35444 |
| TPR KD | LB2 | 125.2 | 54.8 | 124.1 | 90.0 | 1.1 | -35.1 | 0.00 |  | -13.6 | 0.00 | 36974 |
| NUP153 KD | LB2 | 139.7 | 60.4 | 129.1 | 64.1 | 10.6 | -3.7 | 0.00 |  | 0.9 | 0.00 | 31628 |
| ELYS KD | LB2 | 120.6 | 56.4 | 126.5 | 62.4 | -5.9 | -6.0 | 0.00 |  | -18.2 | 0.00 | 25215 |

Median and standard deviation of the observed and expected lamin face to NPC distances, the difference between them, p-values (see Methods), and number of NPCs. The distributions were also compared to scrambled siRNA control. The data in each row was collected from 20 cells. The Mann Whitney U test and Ansari-Bradley test were used as in described in the Materials and Methods.

**Table 2C:** Face radii distributions (fiber to NPC + face to NPC) of WT MEFs with TPR, NUP153, and ELYS knockdown.

| siRNA<br>Knockdown | Lamin<br>Labeled | Observed (nm) |  | Expected (nm) |  | Obs. - Exp. (nm) |  | P vs Exp. |  | Obs. - Scram.<br>Median (nm) | P vs Scram.<br>Median | Num. of NPCs<br>N |
| --- | --- | --- | --- | --- | --- | --- | --- | --- | --- | --- | --- | --- |
|  |  | Median | St. Dev. | Median | St. Dev. | Median | St. Dev. | Median | St. Dev. |  |  |  |
| Scrambled | LA | 163.3 | 53.2 | 171.9 | 67.4 | -8.6 | -14.2 | 0.00 |  |  |  | 39096 |
| TPR KD | LA | 154.3 | 49.6 | 164.3 | 59.9 | -10.0 | -10.4 | 0.00 |  | -9.1 | 0.00 | 40767 |
| NUP153 KD | LA | 155.9 | 48.3 | 162.8 | 56.6 | -6.9 | -8.3 | 0.00 |  | -7.5 | 0.00 | 36066 |
| ELYS KD | LA | 169.7 | 50.9 | 168.9 | 72.3 | 0.8 | -21.3 | 0.38 | 0.00 | 6.3 | 0.00 | 21521 |
| Scrambled | LC | 157.0 | 50.8 | 163.3 | 77.4 | -6.4 | -26.6 | 0.00 |  |  |  | 37760 |
| TPR KD | LC | 150.8 | 47.0 | 161.5 | 103.3 | -10.7 | -56.2 | 0.00 |  | -6.2 | 0.00 | 35489 |
| NUP153 KD | LC | 152.8 | 47.3 | 157.8 | 58.9 | -4.9 | -11.7 | 0.00 |  | -4.1 | 0.00 | 39988 |
| ELYS KD | LC | 167.5 | 52.0 | 169.0 | 77.1 | -1.5 | -25.1 | 0.00 |  | 10.5 | 0.00 | 27053 |
| Scrambled | LB1 | 174.7 | 54.7 | 181.8 | 92.2 | -7.1 | -37.5 | 0.00 |  |  |  | 37383 |
| TPR KD | LB1 | 154.4 | 48.0 | 163.2 | 89.4 | -8.9 | -41.4 | 0.00 |  | -20.3 | 0.00 | 40899 |
| NUP153 KD | LB1 | 173.6 | 56.1 | 178.1 | 67.1 | -4.4 | -11.0 | 0.00 |  | -1.1 | 0.06 | 31145 |
| ELYS KD | LB1 | 157.1 | 48.8 | 162.1 | 70.6 | -5.0 | -21.7 | 0.00 |  | -17.6 | 0.00 | 24981 |
| Scrambled | LB2 | 175.5 | 52.5 | 175.0 | 129.5 | 0.4 | -76.9 | 0.22 | 0.95 |  |  | 35444 |
| TPR KD | LB2 | 159.0 | 47.7 | 158.7 | 147.2 | 0.3 | -99.4 | 0.16 | 0.40 | -16.4 | 0.00 | 36974 |
| NUP153 KD | LB2 | 170.6 | 55.2 | 166.5 | 68.9 | 4.0 | -13.7 | 0.00 |  | -4.9 | 0.00 | 31628 |
| ELYS KD | LB2 | 160.7 | 48.7 | 162.7 | 70.3 | -2.0 | -21.6 | 0.00 |  | -14.8 | 0.00 | 25215 |

Median and standard deviation of the observed and expected sum of lamin fiber and lamin face to NPC distances, the difference between them, p-values (see Methods), and number of NPCs. The distributions were also compared to scrambled siRNA control. The data in each row was collected from 20 cells. p-values less than the Bonferroni corrected alpha value are in red. The Mann Whitney U test and Ansari-Bradley test were used as in described in the Materials and Methods.

**Figure 5.**
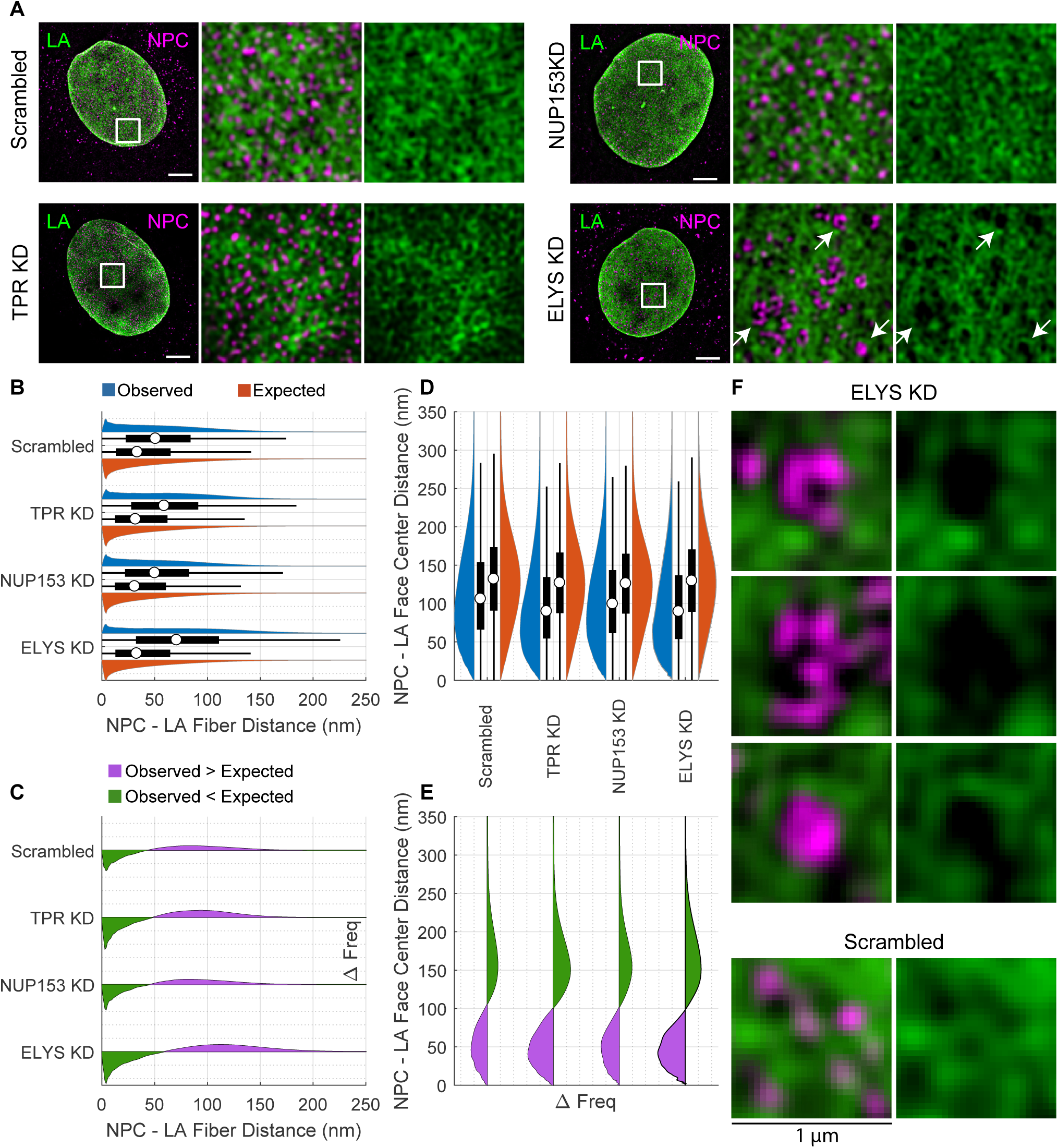
Co-distribution of LA and NPC components after siRNA transfection show enlarged LA meshworks filled with NPC clusters upon ELYS knockdown. A) Immunofluorescence images of LA (green) and NPCs (magenta) following knockdowns (KD) of TPR, NUP153, ELYS and scramble control. Note the clustering of NPCs in the ELYS KD. Area of white box (left) is shown merged (center) and just lamin (right). White arrows indicate areas of NPC clustering. Scale bar = 5 *µm*. B) Paired violin and box plots of NPC center to LA fiber center distances. The violin (blue) and box plots represent the observed distance distributions. The violin (red) and box plots on bottom represent the expected distance distributions under the null hypothesis. The white circle indicates the median. The thick black bar indicates the interquartile range (IQR). The black whiskers indicate 1.5 times the IQR. C) Frequency difference plots of observed minus expected LA fiber to NPC distances for the knockdown series. The green portion below the line indicates where the observed frequency is less than expected. The purple portion above the line indicates where the observed frequency is greater than expected. D) NPC center to LA face center distances displayed as in (B), rotated 90 degrees counterclockwise. E) Frequency difference plot of NPC to LA face center distances, displayed as in (C), rotated 90 degrees counterclockwise.F) 1 *µm*^2^ areas around NPC clusters formed after scramble treatment or ELYS KD indicated by white arrows in (A) shown merged (left) and just lamin (right).

While there did not appear to be NPC clustering upon TPR depletion, the NPCs appeared to be less associated with the LA fibers and more centered within the faces of a dense LA meshwork (Figure 5A). The median distance between the centers of NPCs and LA fibers with TPR knockdown (59.0 nm; Table 2A, Figure 5 B,C, Figure S6) increased versus a scrambled siRNA control, though to a lesser magnitude than for ELYS knockdown (+8.2 nm TPR KD vs +20.0 nm ELYS KD; p < 0.001; Table 2A, Figure 5 B,C). The median distance between NPCs and LA face centers (90.0 nm; Table 2B, Figure 5D) was reduced with TPR knockdown (−16.2 nm; p < 0.001; Table 2B, Figure 5 D, E). The face radius of the LA fiber meshwork (154.3 nm; p < 0.001; Table 2C) was decreased upon TPR depletion (−9.1 nm; p < 0.001; Table 2C). These data suggested that the NPCs were less closely associated with LA fibers following TPR knockdown. Additionally, the reduced face size suggested that the LA meshwork faces were reduced in size (e.g., compacted) upon TPR knockdown forcing NPCs into more confined spaces than in WT LA meshworks.

In contrast to ELYS and TPR knockdowns, NUP153 knockdown only slightly reduced the median distance between NPCs and LA fibers (−0.8 nm; p < 0.001; Table 2A, Figure 5B, C). This reduction was an order of magnitude smaller than observed for the knockdown of either ELYS or TPR. The distance between LA face centers and NPCs was reduced (−6.5 nm; p < 0.001; Table 2B, Figure 5 D, E, Figure S6) and the face radius for the LA meshwork was reduced (−7.5 nm; p < 0.001; Table 2C). The faces in the LA meshwork appeared smaller and more compact compared to controls which was similar to the effect seen with TPR knockdown. Thus, upon NUP153 knockdown, the faces in the LA meshwork became smaller compared to the scramble control, modestly decreasing both the LA fiber-NPC and LA face-NPC distances. The effect of NUP153 knockdown is similar to that of TPR knockdown but reduced in magnitude.

### Changes in LC meshworks are similar to LA meshworks but of lesser magnitude following silencing of ELYS, TPR and NUP153

Our analysis of LC fibers and NPCs suggested that LC fibers do not have a definable relationship with NPCs in WT MEFs (see Figure 3). However, the co-distribution of LC fibers and NPCs was significantly modified by knockdown of either ELYS or TPR. ELYS knockdown resulted in an increase in the median distance between NPCs and LC fibers (63.1 nm; +20.2 nm vs scrambled; p < 0.001; Table 2A, Figure 6 A,B,C, Figure S7) and the LC face center to NPC center distances decreased (96.1 nm; −13.0 nm vs scrambled; p < 0.001; Table 2B, Figure 6 D,E,F). The knockdown of ELYS also increased the effective face radius (167.5 nm; +10.5 nm vs scrambled; p < 0.001; Table 2C) indicating that ELYS knockdown results in expanded LC meshworks as it did for LA meshworks. These results suggest that the NPC clusters induced by ELYS depletion exclude LC fibers as well as LA fibers.

**Figure 6.**
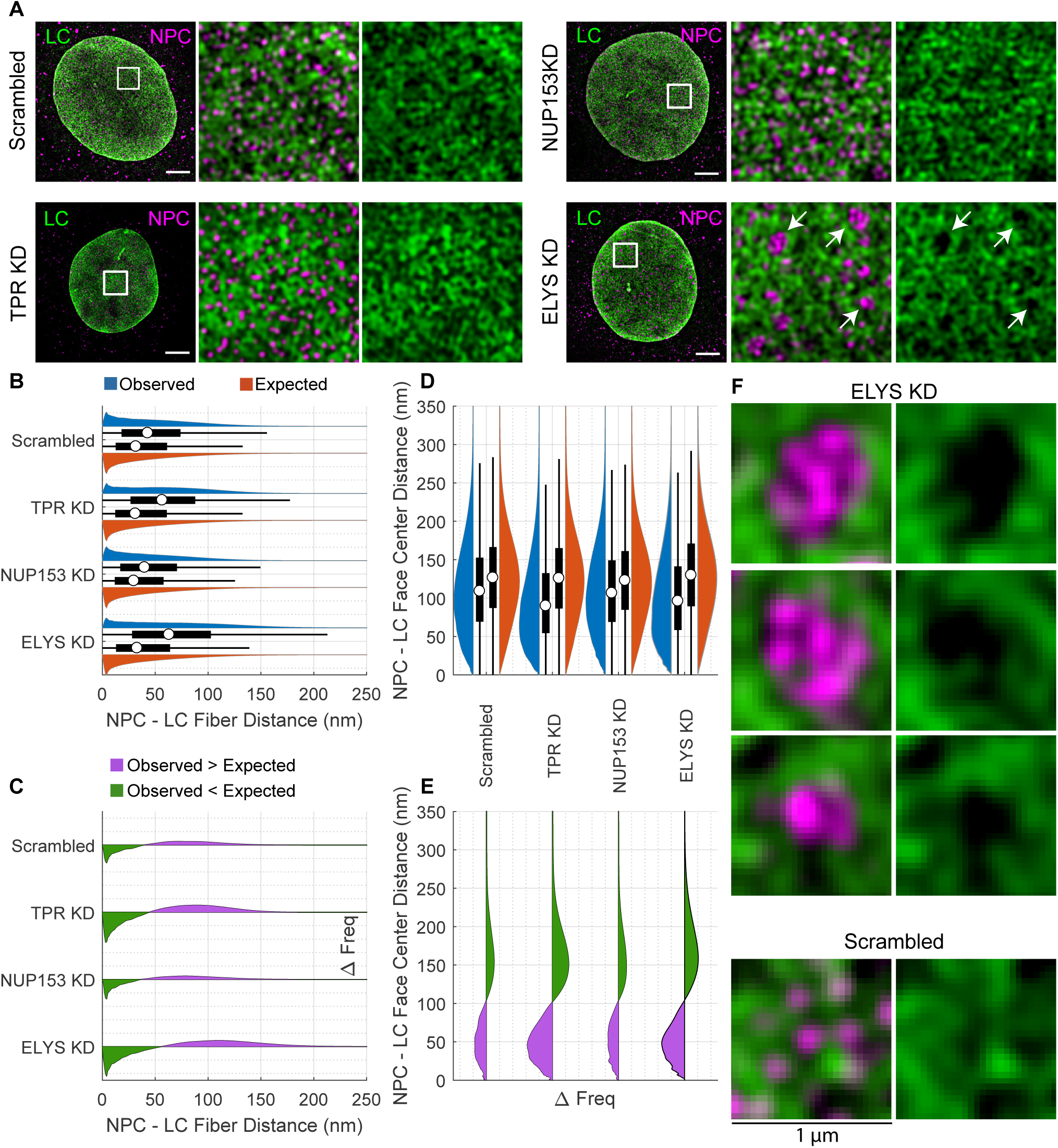
Co-distribution of LC and NPC components after siRNA transfection show enlarged LC meshwork filled with NPC clusters upon ELYS knockdown. A) Double label immunofluoresence images of LC (green) and NPCs (magenta) following KDs of TPR, NUP153, ELYS and scramble control. Area of white box (left) is shown merged (center) and just lamin (right). White arrows indicate areas of NPC clustering. Scale bar = 5 *µm*. B. Paired violin and box plots of NPC center to LC fiber center distances. The violin (blue) and box plots on top represent the observed distance distributions. The violin (red) and box plots on bottom represent the expected distance distributions under the null hypothesis. The white circle indicates the median. The thick black bar indicates the interquartile range (IQR). The black whiskers indicate 1.5 times the IQR. C) Frequency difference plots of observed minus expected LC fiber to NPC distances for the knockdown series. The green portion below the line indicates where the observed frequency is less than expected. The purple portion above the line indicates where the observed frequency is greater than expected. D) NPC center to LC face center distances displayed as in (B), rotated 90 degrees counterclockwise. E) Frequency difference plot of NPC center to LC face center distances, displayed as in (C), rotated 90 degrees counterclockwise. F) 1 *µm*^2^ areas around NPC clusters formed after scramble treatment or ELYS KD indicated by white arrows in (A) shown merged (left) and just lamin (right).

siRNA knockdown of TPR resulted in an increase in the median distance between NPCs and LC fibers (+13.7 nm vs scramble; p < 0.001; Table 2A, Figure 6B, C, Figure S7), a decrease in median distances between NPCs and LC face centers (−19.2 nm; p < 0.001; Table 2B, Figure 6 D,E) and a decrease in the effective face radius (−6.2 nm; Table 2C; p < 0.001). These results indicate that the LC meshwork face size decreased after TPR knockdown, similar to LA.

NUP153 knockdown resulted in a decrease (−3.0 nm; p < 0.001; Table 2A, Figure 6 B, C, Figure S7) in the median distance between NPCs and LC fibers. Decreases in LC face to NPC center distances (−2.2 nm; p < 0.0.01; Table 2B, Figure 6 D,E) and face radius were also detected (−4.1 nm; p < 0.001; Table 2C). While these decreases are consistent with the change seen in the distances between NPCs and LA fibers, the magnitude of the change is much less than for depletion of ELYS or TPR. Overall, the observed changes in the NPC distribution relative to LC fibers upon ELYS, TPR, and NUP153 knockdown were similar to those observed for LA fibers.

### Depletion of TPR or NUP153 results in denser LB1 meshworks, while LB1 fibers protrude into NPC clusters upon ELYS knockdown

Depletion of TPR, NUP153, or ELYS altered the median center-to-center distance between LB1 fibers and NPCs (+0.5 nm, −4.7 nm, and −3.1 nm, respectively, Obs. – Scram; p < 0.001; Table 2A, Figure 7A, B, Figure S8) relative to scrambled siRNA controls. The small magnitude of these changes suggests that depletion of these nucleoporins had a minimal impact on the relationship between LB1 and NPCs compared to the changes seen in the distances between NPCs and LA/C fibers (Figure 7C). In contrast, the changes in median distance between LB1 face centers and NPCs were larger in magnitude upon knockdown of TPR, NUP153, or ELYS (−19.2 nm, −2.5 nm, and −13.0 nm, respectively; Obs. – Scram.; p < 0.001; Table 2B, Figure 7 D, E, F, Figure S8) ; and face radii decreased (−20.3 nm, −1.1 nm, −-17.6 nm; Obs. – Scram.; p < 0.001; Table 2C). Knocking down TPR or ELYS decreased the distances between NPCs and LB1 face centers as well as the LB1 face radii, while knocking down NUP153 had less impact.

**Figure 7.**
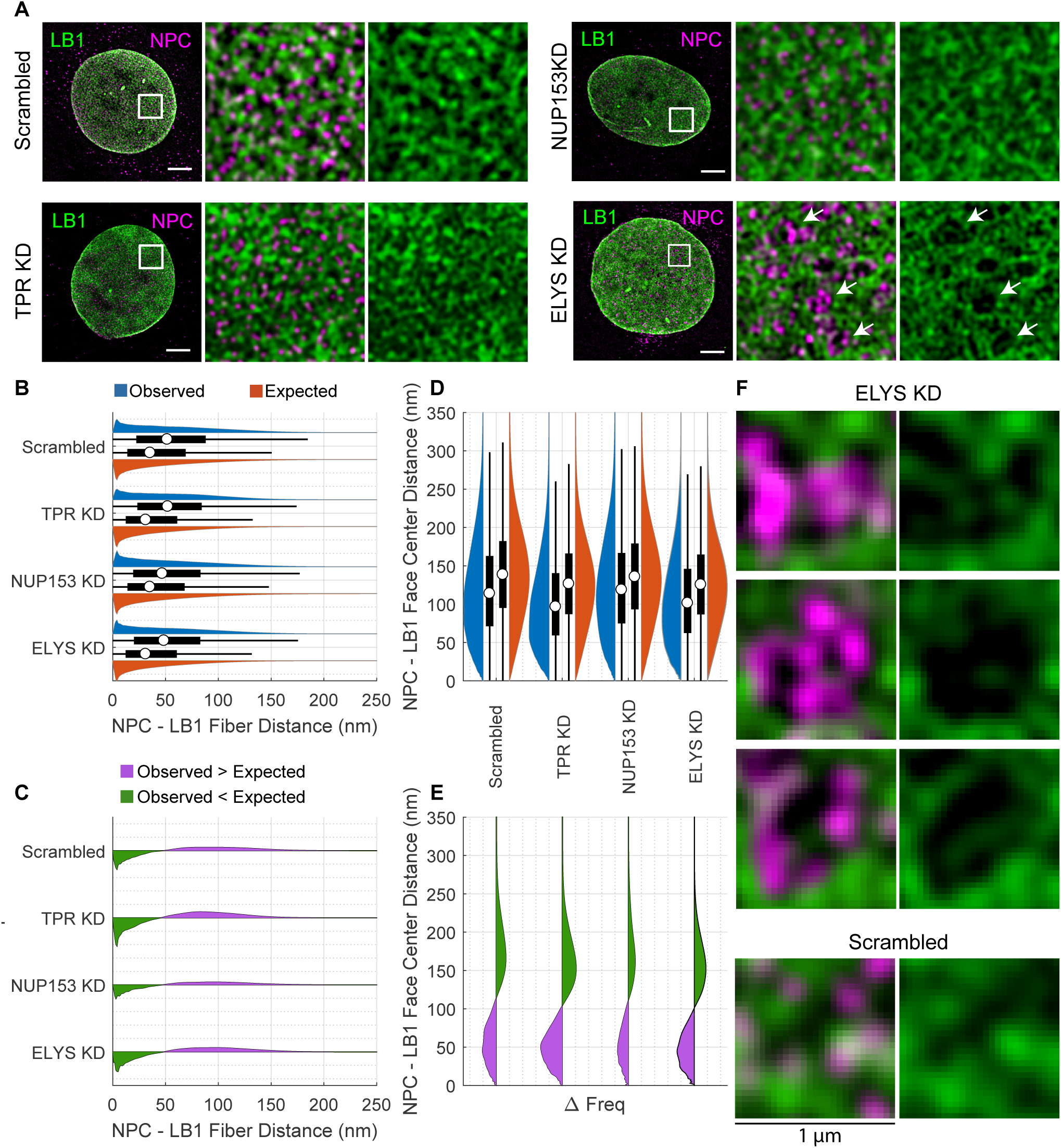
Co-distribution of LB1 and NPCs after siRNA transfection reveal LB1 fibers within NPC clusters upon ELYS knockdown. A) Double label immunofluoresence images of LB1 (green) and NPCs (magenta) following KDs of TPR, NUP153, ELYS and scramble control. Area of white box (left) is shown merged (center) and just lamin (right). White arrows indicate areas of NPC clustering. Scale bar = 5 *µm*. B) Paired violin and box plots of NPC center to LB1 fiber center distances. The violin (blue) and box plots on top represent the observed distance distributions. The violin (red) and box plots on bottom represent the expected distance distributions under the null hypothesis. The white circle indicates the median. The thick black bar indicates the interquartile range (IQR). The black whiskers indicate 1.5 times the IQR. C) Frequency difference plot of observed minus expected LB1 fiber to NPC center distances for the knockdown series. The green portion below the line indicates where the observed frequency is less than expected. The purple portion above the line indicates where the observed frequency is greater than expected. D) NPC center to LB1 face center distances displayed as in (B), rotated 90 degrees counterclockwise. E) Frequency difference plot of NPC to LB1 face center distances, displayed as in (C), rotated 90 degrees counterclockwise. F) 1 *µm*^2^ areas around NPC clusters formed after scramble treatment or ELYS KD indicated by white arrows in (A) shown merged (left) and just lamin (right).

Visual inspection of the accompanying images reveals denser LB1 meshworks upon TPR and NUP153 depletion relative to scrambled siRNA controls as the numerical analysis suggests, but also enlarged faces upon ELYS knockdown in contrast with the quantitative measurements. Closer inspection of the images upon ELYS depletion reveals LB1 fibers protruding into the enlarged faces (Figure 7). This is not seen in the enlarged faces of LA/C meshworks (Figure 5A and 6A). The interdigitation of LB1 fibers within the NPC clusters explains why an increase in LB1 fiber to NPC distances is not seen quantitatively.

### Depletion of ELYS, TPR, or Nup153 has a minor impact on the independence between LB2 fibers and NPCs

As described in previous sections, we could not detect a relationship between LB2 fibers and NPCs in WT MEFs (see Figure 3). Upon knockdown of TPR, NUP153, or ELYS, the observed distances between LB2 fibers and NPCs differed by a few nanometers from expected (−1.7 nm, −6.6 nm, and +3.0 nm, respectively; Obs.- Exp., p < 0.01; Table 2A, Figure 8A,B, Figure S9) and from the scramble control (−1.5 nm, −4.4 nm, and +4.1 nm, respectively; Obs. - Scram; p < 0.01; Table 2A, Figure 8A,B,C). Although the changes in association between the NPCs and LB2 fibers were minimal, the differences were statistically significant with NUP153 knockdown having the greatest effect. In contrast, LB2 face center to NPC center distances (−13.6 nm, +0.9 nm, and −18.2 nm vs scrambled; Obs. – Scram.; p < 0.01; Table 2B; Figure 8D,E,F) and the face radii decreased significantly (−16.4 nm, −4.9 nm, −14.8 nm vs scrambled; Obs. – Scram; p < 0.01; Table 2C,, Figure S9), following knockdown of TPR, NUP153, or ELYS, respectively. Thus, the main effect of the TPR and ELYS knockdown was to decrease the LB2 face radii and the distance to the LB2 face centers relative to the NPC distribution. In contrast, the LB2 fiber to NPC center distances were not perturbed to the same extent when compared to the other lamin fibers.

**Figure 8.**
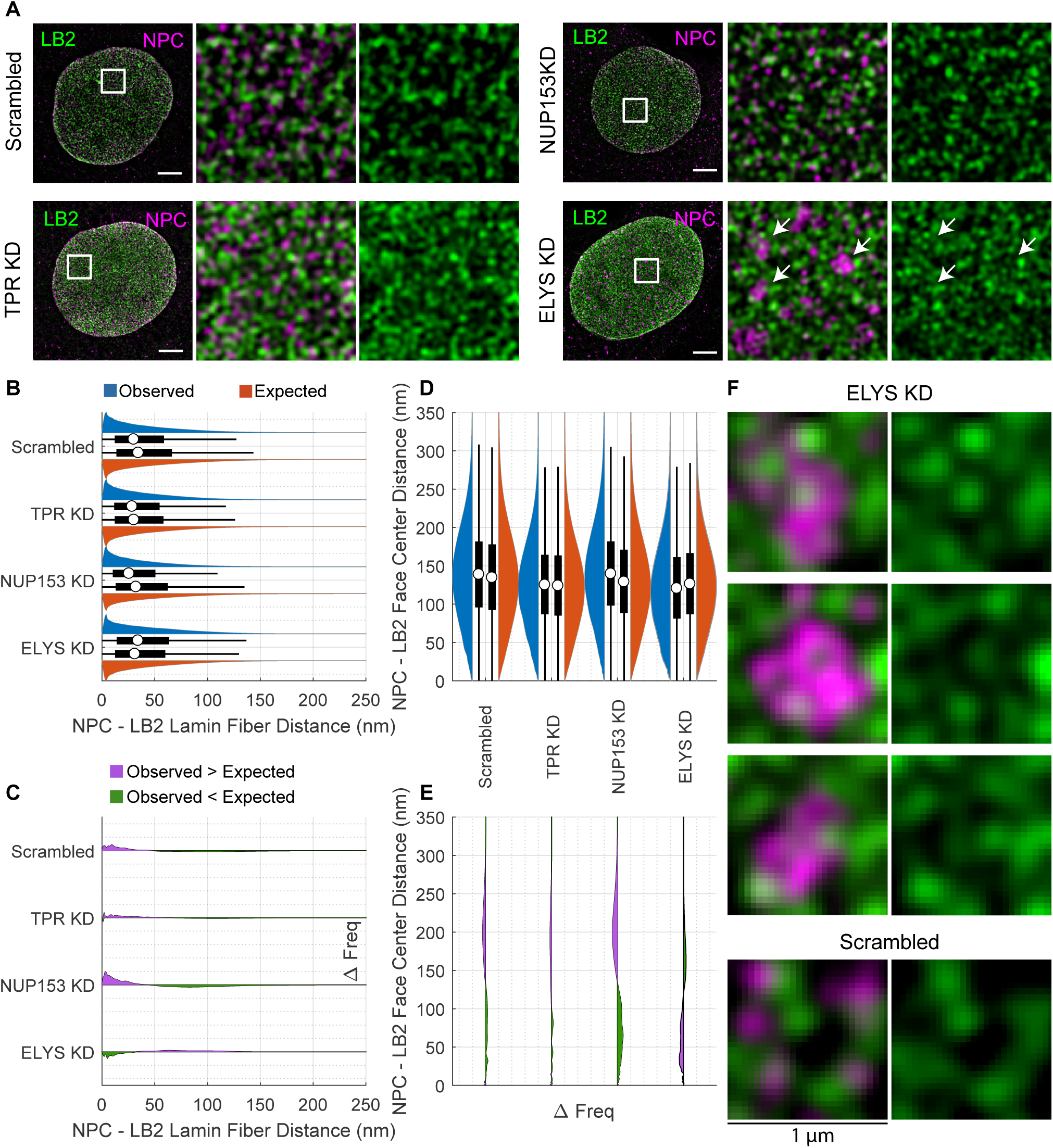
Co-distribution of LB2 and NPCs after siRNA transfection do not show enlarged faces around NPC clusters upon ELYS knockdown. A) Immunofluorescence images of LB2 (green) and NPCs (magenta) following KDs of. TPR, NUP153, ELYS and scramble control. Area of white box (left) is shown merged (center) and just lamin (right). White arrows indicate areas of NPC clustering. Scale bar = 5 µm. B) Paired violin and box plots of NPC center to LB2 fiber center distances. The violin (blue) and box plots on top represent the observed distance distributions. The violin (red) and box plots on bottom represent the expected distance distributions under the null hypothesis. The white circle indicates the median. The thick black bar indicates the interquartile range (IQR). The black whiskers indicate 1.5 times the IQR. C) Frequency difference plot of observed minus expected LB2 fiber center to NPC center distances. The green portion below the line indicates where the observed frequency is less than expected. The purple portion above the line indicates where the observed frequency is greater than expected. D) NPC center to LB2 face center distances displayed as in (B), rotated 90 degrees counterclockwise. E) Frequency difference plot of NPC to LB2 face center distances, displayed as in (C), rotated 90 degrees counterclockwise. F) 1 *µm*^2^ areas around NPC clusters formed after scramble treatment or ELYS KD indicated by white arrows in (A) shown merged (left) and just lamin (right).

### Silencing of nucleoporins have distinct clustering effects in *Lmna*^*-/-*^ and *Lmnb1*^*-/-*^ MEFs

In addition to the NPC clustering following ELYS knockdown in WT MEFs (Figure 5A), we observed similar NPC clustering following ELYS knockdown in *Lmna*^*-/-*^ and *Lmnb1*^*-/-*^ MEFs (Figure S10A). This suggest the clustering effect induced by ELYS depletion is not strongly dependent on the presence of LA/C or LB1.

NUP153 knockdown had modest effects on the relationship of NPC to lamin fiber distances and lamin meshwork sizes in WT cells. However, we did observe clustering of NPCs in *Lmna*^*-/-*^ and *Lmnb1*^*-/-*^ upon knockdown of NUP153 (Figure S10B).

With TPR knockdown, we did not see an increase in the number of NPCs or clustering compared to scrambled siRNA in WT MEFs (Figure S10C,D). The only clear change in the number of NPCs in WT MEFs was upon ELYS KD, but this may be due to our inability to resolve individual NPCs in the the clusters that formed (p < 0.001).

Across the ten cells analyzed, the change in the median number of NPCs observed in *Lmnb1*^*-/-*^ MEFs was not significant upon TPR KD versus scramble control (Figure S10D). However, the shape of the distribution of the number of NPCs following TPR knockdown did appear altered in *Lmnb1*^*-/-*^ MEFs with a sub-population showing a similar number of NPCs as WT MEFs. This leaves open the possibility that effects on the number of NPCs following TPR KD may be dependent on the amount of LB1 present in the cell (Figure S10D).

## Discussion

Ever since the first descriptions of the NE as a distinct structure in eukaryotic cells, the relationships between the components of the structure have been the subject of intense scrutiny. However, due to multiple factors, including its dense composition, relative insolubility and thin structure sandwiched between the chromatin and the cytoplasm, determination of its fine structure has been elusive. Several lines of evidence support the consensus that NPCs are anchored to the lamina during interphase. Studies of the dynamics of both lamins and NPCs in interphase cells show that neither has appreciable lateral mobility in the NE (Moir et al., 2000; Daigle et al., 2001). Biochemical fractionation of the NE as well as electron microscopy studies of both somatic cells and amphibian eggs demonstrated that lamins and NPCs are intimately associated (Dwyer and Blobel, 1976; Gerace et al., 1984; Scheer et al., 1976).

Our 3D-SIM imaging and quantitative analysis of the MEF nuclei constitute a data set that reveals important insights into the structural relationship between the lamin fibers and NPCs. The image analysis focuses on localizing structures, lamin fibers, and NPCs, to high precision and then performing statistical analysis on the aggregate data set. This is distinct from localizing individual fluorophores through single molecule localization microscopy (SMLM), the Delaunay triangulation (DT) of those fluorophore localizations, or subgraphs of the DT such as the Euclidean Minimum Spanning Tree (Xie et al., 2016; Kittisopikul et al., 2019). Extracting information about fibrous lamin structures from SMLM data would require additional analysis not directly realizable from SMLM localizations or their graphs (Peters et al., 2018; Kittisopikul et al., 2019). Our analysis of lamin fibers as employed here has been purpose built and validated for use in dense structures such as lamin meshworks with complex junctions (Kittisopikul et al., 2020). Electron microscopy as well as the meshwork altering perturbations produced here suggest the fibrous nature of lamins exists even in the dense wild-type lamina. To evaluate the relationship between lamin fibers and NPCs to high precision, we have exploited the continuous nature of the imaging data set afforded by Nyquist sampling to localize structures by mathematical optimization as described in the Appendix. The combination of super-resolution microscopy and computational analysis as a data set will allow researchers to pursue further questions about the relationship of lamin fibers and NPCs as we have demonstrated here.

Which of the lamin isoforms interact with the NPCs has been a relevant question, since the four major lamin isoforms, LA, LC, LB1, and LB2 are not all expressed throughout development and each may not be expressed in all cell types (Burke and Stewart, 2014). With the aid of super resolution microscopy techniques, it is now established that each of the lamin isoforms assembles into a distinct network in the NE (Shimi et al., 2015; Xie et al., 2016) and the relationship of NPCs with each lamin isoform can be determined with increasing precision. Studies on the cell cycle dependent dynamics of NPCs have identified so-called ‘pore-free islands’ in G1 nuclei of multiple cell types (Maeshima et al., 2006; Mimura et al., 2017). These pore-free areas are enriched in LA/C and generally devoid of B-type lamins. Ectopic over expression of LA could induce the formation of pore-free islands while depletion of LA/C by siRNA knockdown dispersed pore-free islands leading to a more uniform distribution of NPCs (Maeshima et al., 2006). The expression of any of several laminopathic forms of LA/C or depletion of LB1 leads to the formation of herniations or blebs in the NE that contain an expanded LA/C meshwork and are generally depleted in B-type lamins (Goldman et al., 2004; Shimi et al., 2008; Mounkes et al., 2003; Raharjo et al., 2001). These blebs are also deficient in NPCs. Together these studies suggest that B-type lamins may be more important than LA/C for the normal distribution of NPCs in the NE. This conclusion makes sense intuitively since stem cells and some differentiated cells express very little or no LA/C, yet have what appears to be a regular distribution of NPCs Burke and Stewart (2014). However, other studies have suggested that lamin isoforms can function redundantly to ensure normal NPC distribution (Guo et al., 2014). Our findings presented here support the notion that both LA and LB1 have clear spatial relationships with NPCs and these relationships are preserved when either LA/C or LB1 is absent. Although the proximal lateral relationship between NPCs and LA and LB1 fibers is retained in both types of lamin null cells, the quantitative data suggest that the presence of LA fibers may be more important to the LB1-NPC relationship than the presence of LB1 fibers are to the LA-NPC relationship. Using cryo-ET, we were able to demonstrate that both LA and LB1 fibers lie in close proximity to the NPC, and in several cases can be seen in intimate association with the nucleoplasmic ring structure of the NPC. This finding supports our super resolution results that indicate a close physical relationship for both LA and LB1 with NPCs over the entire nucleus. Measurement of LC interactions with NPCs followed a similar trend to those of LA and LB1 in our analyses, although we could not draw firm conclusions on the LC-NPC interaction due to the small magnitude of the observed values relative to expected. Surprisingly, we did not find an obvious relationship between LB2 and NPCs in our analysis.

Xie and coworkers have previously carried out super resolution microscopy studies of the relationships between lamins and NPCs in mouse adult fibroblasts (MAFs) (Xie et al., 2016). By re-expressing mEOS-tagged LA or LC in Lmna-/- cells, they found NPCs concentrated in the spaces between LA fibers, and a close association of NPCs with the LC networks. These findings are directly the opposite of those we report here. There are several possible explanations for these discrepancies including: 1) possible differences between adult fibroblasts and embryonic fibroblasts, 2) possible differences in an ectopically expressed lamin network versus the endogenous networks, and 3) over-expression of LA only or LC only versus cells expressing all four lamin isoforms in the natural ratio. Further studies will be necessary to address these differences in results.

Our results also provide new and important insights into lamin-NPC interaction by knocking down specific nucleoporin levels using siRNA for ELYS, TPR, or NUP153. Each knockdown had unique effects on both NPC distribution and lamin meshwork structure. ELYS knockdown caused dramatic changes in NPC distribution attributable to NPCs clustering within the open faces formed by all of the lamin meshworks and a reduction in NPC number. Depletion of ELYS also led to an increase in the lamin fiber to NPC distance for LA, LC and LB2, but a decrease in the LB1 to NPC distance. NPCs form in a biphasic pattern; at the end of mitosis as the NE reforms and then during interphase (Doucet et al., 2010). These two processes differ in the order that nucleoporins assemble and the enzymatic requirements for assembly. The postmitotic phase involves the recruitment of the NUP107-160 subcomplex to the chromatin surface by the binding of one component of the complex, ELYS/MEL-28 to nucleosomes (Rasala et al., 2006; Galy et al., 2006; Gómez-Saldivar et al., 2016). While we have not demonstrated a direct interaction between ELYS and the lamins, it is clear that the presence of ELYS is required to maintain lamin-NPC interactions. The clustering of NPCs after ELYS knockdown is likely due to the failure of NPCs to correctly assembly on chromatin following mitosis suggesting that, at least for NPCs formed at NE reformation, their association with lamins occurs at that time. ELYS knockdown has previously been shown to decrease NPC density (Doucet et al., 2010; Jevtić et al., 2019; Mimura et al., 2016), disrupt the proper localization of the integral inner nuclear membrane protein lamin B receptor (LBR) (Mimura et al., 2016), and cause the cytoplasmic accumulation of LB1 (Jevtić et al., 2019; Mimura et al., 2016). However, these previous studies did not find clustering of NPCs or changes in lamin meshwork structure.

TPR is a nucleoporin located in the nuclear basket structure of the NPC and could act as a negative regulator of NPC number (McCloskey et al., 2018). In contrast two other studies found that siRNA reduction of TPR reduced NPC number (Funasaka et al., 2012; Fišerová et al., 2019). In our experiments, we also observed a small, but statistically significant increase in NPC numbers after TPR knockdown in WT cells. When we depleted TPR in *Lmna*^*-/-*^ and *Lmnb1*^*-/-*^ cells, a similar small increase in NPCs was observed suggesting that neither lamin isoform alone is involved in regulating NPC numbers. As with ELYS knockdown, TPR knockdown resulted in displacement of the NPCs away from the lamin fibers, with the exception of LB2 fibers, which were slightly closer to the NPCs when TPR was depleted. NUP153 depletion had the most consistent effects on the lamin fiber-NPC relationship with a decrease in lamin fiber to NPC distance and a compaction of the lamin meshworks, although these changes were more modest than those of the other nucleoporin knockdowns. Surprisingly, knockdown of NUP153 in Lmna-/- and Lmnb1-/- cells led to clustering of NPCs in the lamin meshwork faces. This suggests that an interaction of NUP153 with both lamin isoforms is required for normal NPC distribution. NUP153 is known to bind to both LA and LB1 (Al-Haboubi et al., 2011).

The results presented here suggest that the lamina structure and NPCs are co-dependent, that is, changing one of the structures has an effect on the other’s distribution. In addition to the NPC clustering in lamin meshwork faces after ELYS reduction, the lamin meshworks became larger for LA and LC, but became smaller for LB1 and LB2. In contrast, knockdown of either TPR or NUP153 caused each of the lamin meshwork faces to decrease in size. Based on these results, it is tempting to speculate that the number of NPCs helps to determine lamin meshwork structure. Our results show that each of the lamin isoforms appears to interact differently with the three nucleoporins. It should be noted that while ELYS is required for post-mitotic NPC assembly (Franz et al., 2007), NUP153 is required for interphase NPC assembly (Vollmer et al., 2015; Franz et al., 2007), whereas TPR is required only for formation of the nuclear basket (Duheron et al., 2014). In cell-free extracts of Xenopus eggs that recapitulate nuclear assembly, the recruitment of Nup153 to the NE is dependent on the formation of the lamina (Smythe et al., 2000). TPR is also required to maintain the heterochromatin exclusion zones found at the NPCs (Krull et al., 2010) and all three nucleoporins are known to affect chromatin modification states (Kuhn and Capelson, 2019). The lamins are also closely associated with chromatin at the nuclear periphery and it is likely that peripheral chromatin is also playing a role in the association of lamins and NPCs and their distribution in the NE.

Overall, the extensive SIM imaging and quantitative analysis performed here provides important biological insight as to how NPCs and lamin fibers are arranged in the mammalian nucleus. In perturbing the cells and their nuclei by either knocking out lamin isoforms, LA/C or LB1, or knocking down nucleoporins, our data set provides knowledge about interactions mediated by those specific lamin isoform and nucleoporins. In particular, it is clear from this data set that knocking down lamin isoforms results in a change in the spatial distribution of NPCs. Additionally, knocking down nucleoporins has an effect on the spatial distribution of the lamin fiber meshwork. Therefore, the lamins and NPCs play a role in organizing each other at the nuclear periphery.

## Materials and Methods

### Sample size estimation

The initial light microscopy images of WT, lamin knockout cells, and the cryo-ET data were acquired before the design of the study and before the computational analysis was developed. Hierarchical power analysis was performed for the siRNA knockdown series of experiments based upon the effect sizes observed in the initial light microscopy images. We sought to evaluate changes in distance between lamin and NPCs as well as changes in NPC number. The limiting factor was the number of cells that needed to be observed in order to detect a ±20% change in number of NPCs per cell with a power of 0.8 at an alpha of 0.01 with the Mann-Whitney U test. The wmwpower package (Mollan et al., 2019) in R (R Core Team, 2016) was used. Using the estimation methods in that package it was determined that imaging 20 cells would exceed those requirements. Based on thousands of distances being measured per cell, it was determined that the power of the lamin-NPC distance studies would also exceed the requirements.

### Replicates

Each experiment was performed in duplicate as technical replicates. Each technical replicate was performed at a distinct time and included all steps from cell culture to fixation and staining. Additionally, for each technical replicate two sets of coverslips were produced. In Tables 1A and 1B, 10 cells were evaluated per row. In Tables 2A, 2B, and 2C, 20 cells were evaluated per row. The cells were distributed across the four coverslips produced. Outliers were not excluded from the data. Microscopy as described below was done on fixed samples in blocks of time using coverslips from multiple technical replicates. Experimental samples and their controls were conducted within the same microscopy session.

### Statistical reporting

Statistical analysis was done in MATLAB (Mathworks, Natick, MA) other than the power calculation down in R as noted above. The frequency of the simulated distances was compared to the observed distances using the Mann-Whitney U test, also known as the Wilcoxon rank sum test. A non-parametric test was used since the Kolmogorov-Smirnov test rejected the null hypothesis that the distributions were normal.

The Mann-Whitney U test evaluated the null hypothesis that the two sets of samples (observed vs expected, ELYS siRNA versus scrambled siRNA, etc) were drawn from the same distribution. If the Mann-Whitney U test failed to reject the null hypothesis for the distance measurements, the Ansari-Bradley test was applied to examine the null hypothesis that the dispersion (i.e. the standard deviation) of the distributions were the same. Bonferroni corrections were applied to the alpha value to compensate for multiple comparisons by dividing an alpha value of 0.05 by the number of comparisons in the table or figure.

### Cell culture

Immortalized WT, *Lmna*^*-/-*^, *Lmnb1*^*-/-*^, and *Lmnb2*^*-/-*^ MEFs were cultured as previously described (Shimi et al., 2015). Briefly, cells were cultured in modified DMEM (Thermo Fisher Scientific, Waltham, MA, USA) supplemented with 10% fetal calf serum, 50 U/ml penicillin G, 50 *µg*/ml streptomycin sulfate (Thermo Fisher Scientific) at 37°C in a humidified CO2 incubator.

### Super resolution microscopy

3D-SIM was carried out as previously described (Shimi et al., 2015). Briefly, a Nikon Structured Illumination Super-resolution Microscope System (Nikon N-SIM; Nikon, Tokyo, Japan) was built on an ECLIPSE Ti-E (Nikon) equipped with a sCMOS camera ORCA-Flash 4.0 (Hamamatsu Photonics Co., Hamamatsu, Japan) and an oil immersion objective lens CFI SR (Apochromat TIRF 100×, NA=1.49, Oil, WD=0.12; Nikon). N-SIM was operated with NIS-Elements AR (Nikon). For image acquisition, 21 optical sections including a region of the lamina were taken at 50-nm intervals. For image reconstruction from the raw data, illumination modulation contrast, high-resolution noise suppression, and out-of-focus blur suppression were set with fixed values of 1, 0.75, and 0.25, respectively. For presentation, images were adjusted for brightness and contrast.

### Indirect immunofluorescence

Samples for indirect immunofluorescence were processed as previously described (Shimi et al., 2015). Cells were seeded on Gold Seal coverglasses (22 × 22 mm2, no. 1.5; Thermo Fisher Scientific) and fixed with methanol for 10 min at −20°C. Lamins were stained with rabbit polyclonal anti-LA (1:500; 323; Dechat et al. (2007)), goat polyclonal anti-LB1 (1:500; SC-6217; Santa Cruz Biotechnology, Dallas, TX, USA), and rabbit monoclonal LB2 (1:100; EPR9701(B); Abcam, Cambridge, MA, USA), and rabbit polyclonal anti-LC (1:500; 321; Dechat et al. (2007)). Nucleoporins were stained with mouse monoclonal MAb414 (1:1000; BioLegend, San Diego, CA). The secondary antibodies used were donkey anti-mouse immunoglobulin G (IgG)–Alexa Fluor 488, donkey anti-mouse IgG–Alexa Fluor 568, donkey anti-rabbit IgG–Alexa Fluor 488, donkey anti-rabbit IgG–Alexa Fluor 568, donkey anti-goat IgG–Alexa Fluor 488, and donkey anti-goat IgG–Alexa Fluor 568 (all 1:500; Thermo Fisher Scientific). Processed coverslips were mounted with ProLong Diamond antifade reagent (Thermo Fisher Scientific).

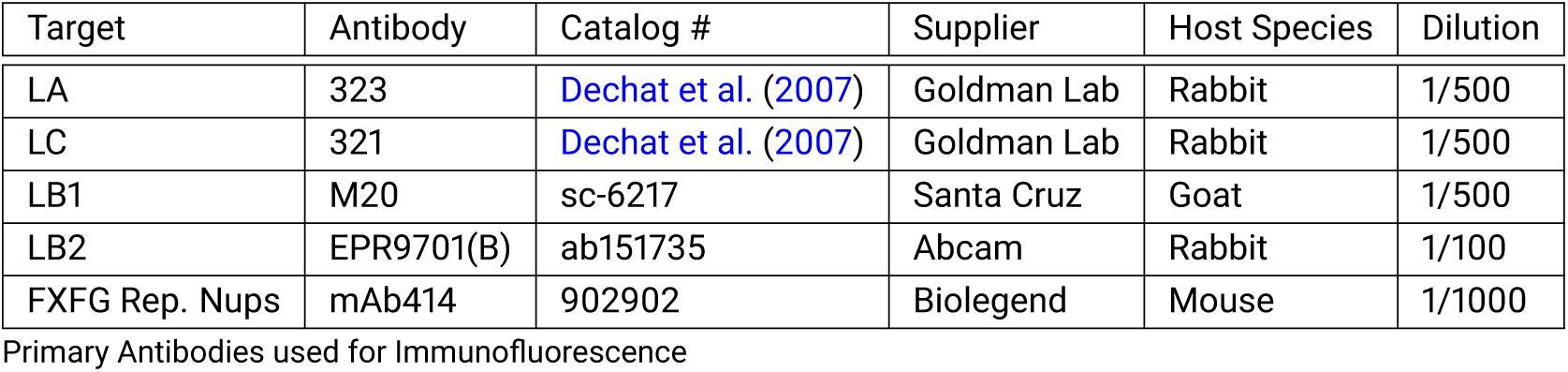

### RNA interference

ON-TARGETplus siRNA oligos (Dharmacon, Lafayette, CO, USA) were used for RNAi-mediated knockdown experiments. Scrambled sequence for control siRNAs;

1. (D-001810-01) 5’-UGGUUUACAUGUCGACUAA-3’
2. (D-001810-02) 5’-UGGUUUACAUGUUGUGUGA-3’
3. (D-001810-03) 5’-UGGUUUACAUGUUUUCUGA-3’
4. (D-001810-04) 5’-UGGUUUACAUGUUUUCCUA-3’

Nup153 siRNAs;

1. (J-057025-11) 5’-CGCUAUGUGCAUUGAUAAA-3’
2. (J-057025-12) 5’-GGGACAGGCUUUGGAGAUA-3’

ELYS siRNA

1. (J-051465-09) 5’-CCACUGAACUAACUACUAA-3’
2. (J-051465-10) 5’-GGAAAGAAGAAGGACGUUA-3’

TPR siRNA;

1. (J-041152-09) 5’- CAACAAACAUUCAUCGGUA-3’
2. (J-041152-10) 5’- CGUGACAUGUACCGAAUUU-3’

5 × 10^4^ MEFs were plated into each well of 6-well plates 24 h before transfection. 30 pmol of siRNA oligos was transfected onto the cells in each well with Lipofectamine RNAiMAX transfection reagents (Thermo Fisher Scientific), following the manufacturer’s instructions. 48h after incubation at 37°C, the transfected cells were trypsinizedand replated at 5 × 10^4^ cells/well into each well of 6-well plates and transfected with 30 pmol of the siRNA. 48h after incubation at 37°C, the transfected cells were trypsinized and replated on coverslips for indirect immunofluorescence or plated into a 60 mm dish for western blotting.

### Quantitative blotting of anti-nucleoporin antibodies

The linearity of antibodies to nucleoporins was determined by immunoblotting of whole cell lysates of WT MEFs. Five samples of MEF lysates containing between 7.5 × 10^3^ to 9 × 10^3^ cells were separated in duplicate lanes of a 7.5% SDS-polyacrylamide gel (SDS-PAGE) and transferred to nitrocellulose for immunoblotting. After transfer, the membrane was briefly rinsed in dH2O and stained with Revert Protein Stain (LI-COR) and imaged in an Odyssey Fc (LI-COR Biosciences, Lincoln NB) at 700nm. The membrane was then washed with Tris-buffered saline (TBS) and blocked in 5% non-fat dry milk (NFM) in TBS for 1hr at room temperature and then in the same solution containing 0.1% Tween 20 for 30 minutes. For incubation with antibodies, the appropriate antibody was diluted in blocking solution with Tween at the indicated concentration (See Table Below) and incubated overnight at 4°C with gentle agitation. The blots were washed 3 times for 5 mins each with TBS containing 0.1% Tween 20. For detection, the appropriate secondary antibodies (Licor IRDye 800CW) were diluted 1:15000 in 5% NFM containing 0.2% Tween 20 and incubated with the membrane for 1hr at room temperature with gentle agitation. The membranes were washed 3X 5 mins each with TBS containing 0.1% Tween 20 and allowed to dry. The dried membranes were imaged in an Odyssey Fc at 800nm.

| Target | Antibody | Catalog # | Supplier | Host Species | Concentration |
| --- | --- | --- | --- | --- | --- |
| Nup153 | R3G1 | sc-101544 | Santa Cruz | Rat | 1 $\mu\text{g mL}^{-1}$ |
| Elys | bs-9880R | bs-9880R | Bioss | Rabbit | 0.1 $\mu\text{g mL}^{-1}$ |
| Tpr | ab84516 | ab84516 | Abcam | Rabbit | 0.2 $\mu\text{g mL}^{-1}$ |

Antibodies used for Western blotting

Images of the total protein stain and specific antibody labeling were analyzed using Empiria Studio Software (LI-COR Biosciences, Lincoln NB). The intensity of the specific antibody labeling in each lane was corrected for protein load using the software and the linearity of the antibody response was determined by the software.

The degree of knockdown for each nucleoporin was determined by SDS-PAGE by loading duplicate samples of each knockdown cell lysate such that the antibody response should be in a linear range, based on the analysis of WT lysates. For quantitation of knockdown, a dilution series of WT lysate was run on the same gel at concentrations that were expected to be in the linear range of the antibody response. After electrophoresis and transfer, the membranes were treated identically to the conditions for determining antibody linearity, imaged in the Odyssey Fc and the images analyzed using Empiria software.

### NPC-lamin rendered view

Cryo-electron tomograms that were acquired previously (Turgay et al., 2017) were further analyzed. The central coordinates of NPCs within cryo-tomograms of NE were determined manually and sub-tomograms (340 nm x 340 nm x 20 nm) were reconstructed in MATLAB, using the TOM toolbox (Nickell et al., 2005). The lamin filaments and NPCs in 4 selected sub-volumes were segmented manually and rendered, using the Amira software package (Thermo Fisher Scientific).

### Immunogold labelling image processing

Sub-tomograms of gold labeled lamins (Turgay et al., 2017) were reconstructed as described above (47 sub-tomograms). The subvolumes containing NPCs (in top-view orientation), were projected along the Z axis, to produce a 2D image. The coordinates of the gold clusters (6 nm and 10 nm) were identified manually and counted. The respective histograms were drawn in Excel (Microsoft).

### Computational image analysis

Computational image analysis was done using MATLAB (Mathworks, Natick, MA) using custom software developed in the Jaqaman Lab. Nikon ND2 files containing image and meta data were loaded into MATLAB using Bioformats (Open Microscopy Environment, Linkert et al. (2010)). Nuclear pore complexes were detected and localized using an adapted point Source Detector routine from the lab of Gaudenz Danuser which involved two-dimensional local maxima detection, Gaussian fitting, and Gaussian mixture modeling. Lamin fibers were segmented using multi-orientation analysis as described in Kittisopikul et al. (2020) to accurately segment a meshwork structure with many junctions. Lamin fibers were further localized as in Appendix 1. The source is available on Github at https://github.com/mkitti/LaminNpcAnalysis

Computation was conducted on Northwestern University’s high performance computing environment, Quest. Files were stored on Northwestern University Research Data Storage Service FSMRESFILES. Globus.org and Box.com were used to transfer files between storage and computational environments.

### Expected distribution of NPCs

In this study, the null hypothesis is that there is no relationship between the position of the lamin fibers and NPCs within the nucleus. To determine if a relationship or an association between lamin fibers and NPCs exists, we used statistical methods to see if the observed distances between lamin fibers and NPCs were significantly different than what would expect under this null hypothesis.

To calculate the expected distribution of NPCs relative to the lamin fibers for each nucleus under that null hypothesis we used a Monte Carlo simulation to randomly place NPCs within the nucleus. 60,000 psuedorandom pairs of numbers representing XY locations of NPCs were selected within the image. If they were not within the mask of the nucleus represented by a complex hull, then the XY locations were rejected. The distance between the remaining XY locations were measured to the nearest lamin fiber location as in Appendix 1. The initial number of pairs was selected empirically such that the distance frequencies would not fluctuate more than 1% for 10 nm bins.

### Image Data Repository

Structured Illumination Microscopy (SIM) data is deposited in the Image Data Repository at https://idr.openmicroscopy.org/ with IDR requisition number XXXXX.

## Acknowledgments

The authors would like to acknowledge the assistance of the Center for Advanced Microscopy and the Nikon Imaging Center at the Feinberg School of Medicine, Northwestern University, for assistance with imaging and the use of the Nikon N-SIM Microscope.

## Additional Information

### Author Contributions

MK, TS, SAA, and RDG conceived of the study. TS and MK performed the light microscopy experiments. MT and OM analyzed Cryo-ET data. MK and SAA ran the Western blots. JRT and YZ provided lamin null cell lines. MK and KJ performed the image and statistical analysis. MK, TS, and MT prepared the figures. All authors contributed to the writing of the paper.

## Appendix 1

### Localization of lamin fibers in orientation space

In order to localize lamin fibers, we use an image analysis algorithm that we previously developed that involves the construction of a three dimensional orientation space by augmenting a 2-D image with orientation as an additional third dimension (Kittisopikul et al., 2020). There we focused on addressing the continuous nature of the orientation dimension, we leave the spatial dimensions discretely sampled and localize line detections to nearest pixel in the Non-Maximum Suppression (NMS) and Non-Local Maxima Suppression (NLMS) procedures.

Here we extend the procedure by using the orientations to localize lines, the lamin fibers, to sub-pixel precision by also treating the spatial dimensions as continuous. Given sufficient signal-to-noise ratios and sampling in excess of that required by the Nyquist-Shannon-Whittaker-Kotelnikov sampling theorem, the spatial dimension could also be treated continuously through interpolation. In particular, we use spline interpolation (Unser, 1999). In that case, we can state the localization problem as solving a system of partial differential equations where ***R***(*x, y, θ*; ***K***) is the steerable filter response at some location (*x, y*) at orientation *θ* at the orientation-resolution ***K***.

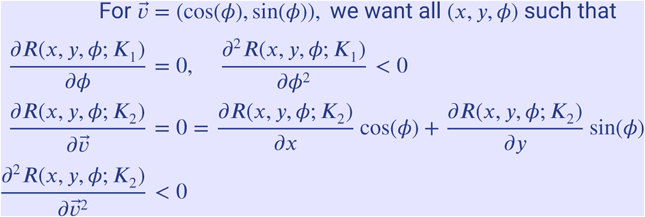

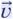 is a vector normal to the structure being localized. As explained in Kittisopikul et al. (2020), ***K***_1_ and ***K***_2_ may differ since the orientation resolution used for orientation detection may differ from the orientation resolution used to localize the detection in space.

### Localization of Lamin Meshwork Face Centers

To understand the relationship of NPCs to the lamin structure, we also measured the distance of the NPCs from their “centers” which we defined as the points furthest away from the lamins within a local neighborhood.

Face centers were localized by identifying local maxima of the distance transform relative to the lamin fibers. A 2D disc with a five pixel radius (150 nm) was used as a structuring element with morphological dilation. This identified the maximum distance within a disc centered at each pixel. The local maxima were detected at the points when the maximum distance within the disc coincided with an identical distance assigned to that pixel via the distance transform. If a connected region with points equidistant from the lamin fibers were found, the centroid of that region was selected as the face center.

Because faces are not always convex or there maybe lamin fibers protruding into faces, multiple distinct centers may be detected. In this case, the distance from the NPC is measured to the nearest face center.

**Figure S1.**
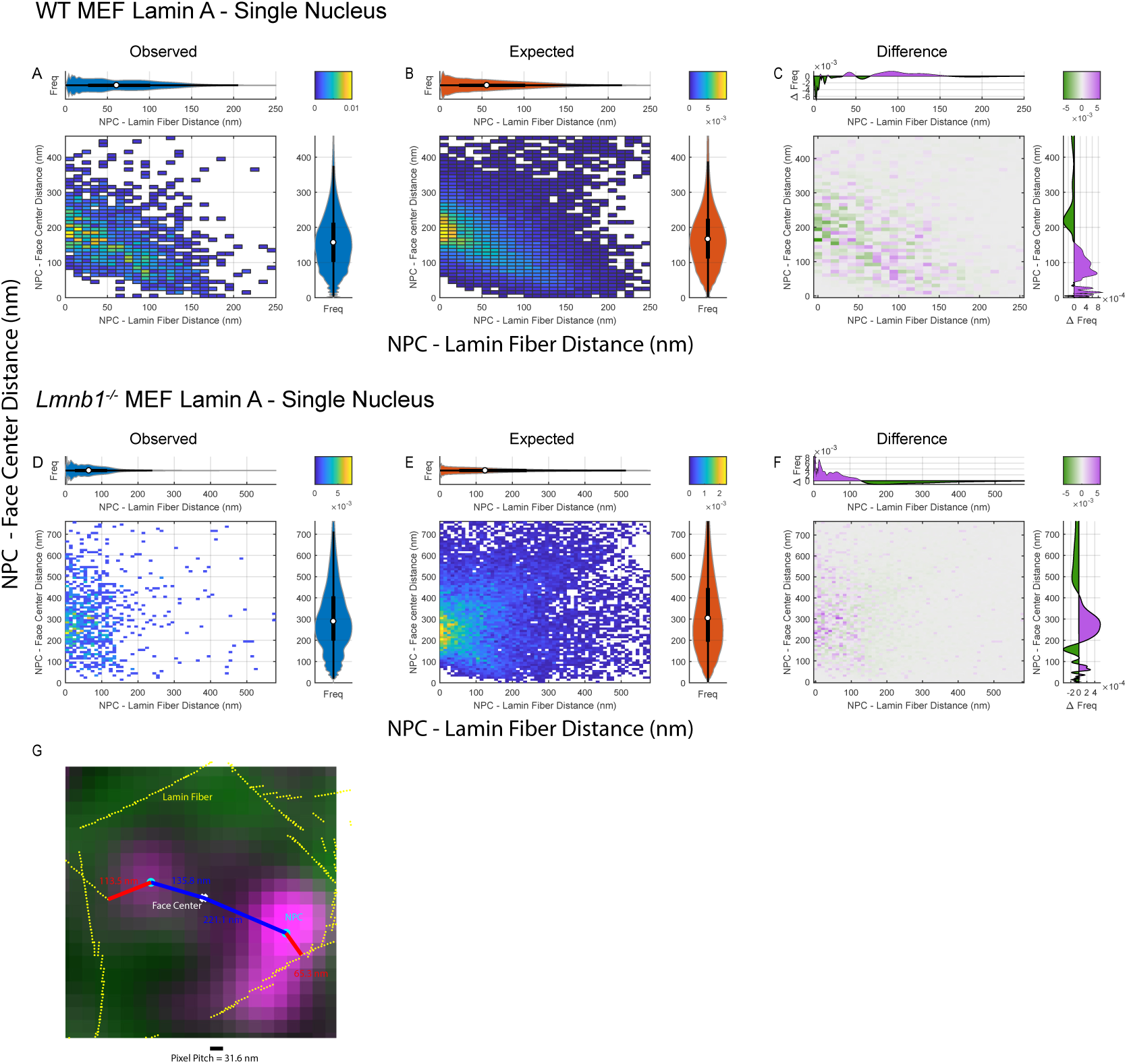
Bivariate histograms of LA fiber-NPC and face center-NPC distances in single nuclei. illustration of distances. A) Observed bivariate histogram of NPC to LA face center distances versus NPC to lamin A fiber distances of a single WT MEF Lamin A nucleus shown in panel A of the Figure 2. B) Expected bivariate histogram of NPC to lamin A face center distances versus NPC to lamin A fiber distances of a single WT MEF Lamin A nucleus under the null hypothesis. C) Difference between the observed and expected distance distributions with purple indicating where the observed exceeds the expected frequency and green showing when the observed frequency is less than the expected frequency. D-F) Same as A-C except for the single Lmnb1-/- nucleus shown in panel A of the Figure 2. Marginal violin plots and box plots of the distances correspond with the half-violin plot counterparts of the same orientation and color as in Panel B of the Figure 2. G) Zoomed in plot showing the NPC to lamin A fiber (red) and NPC to lamin A face center distances (blued) measured. Other colors correspond with those as in panel B of Figure 2.

**Figure S2.**
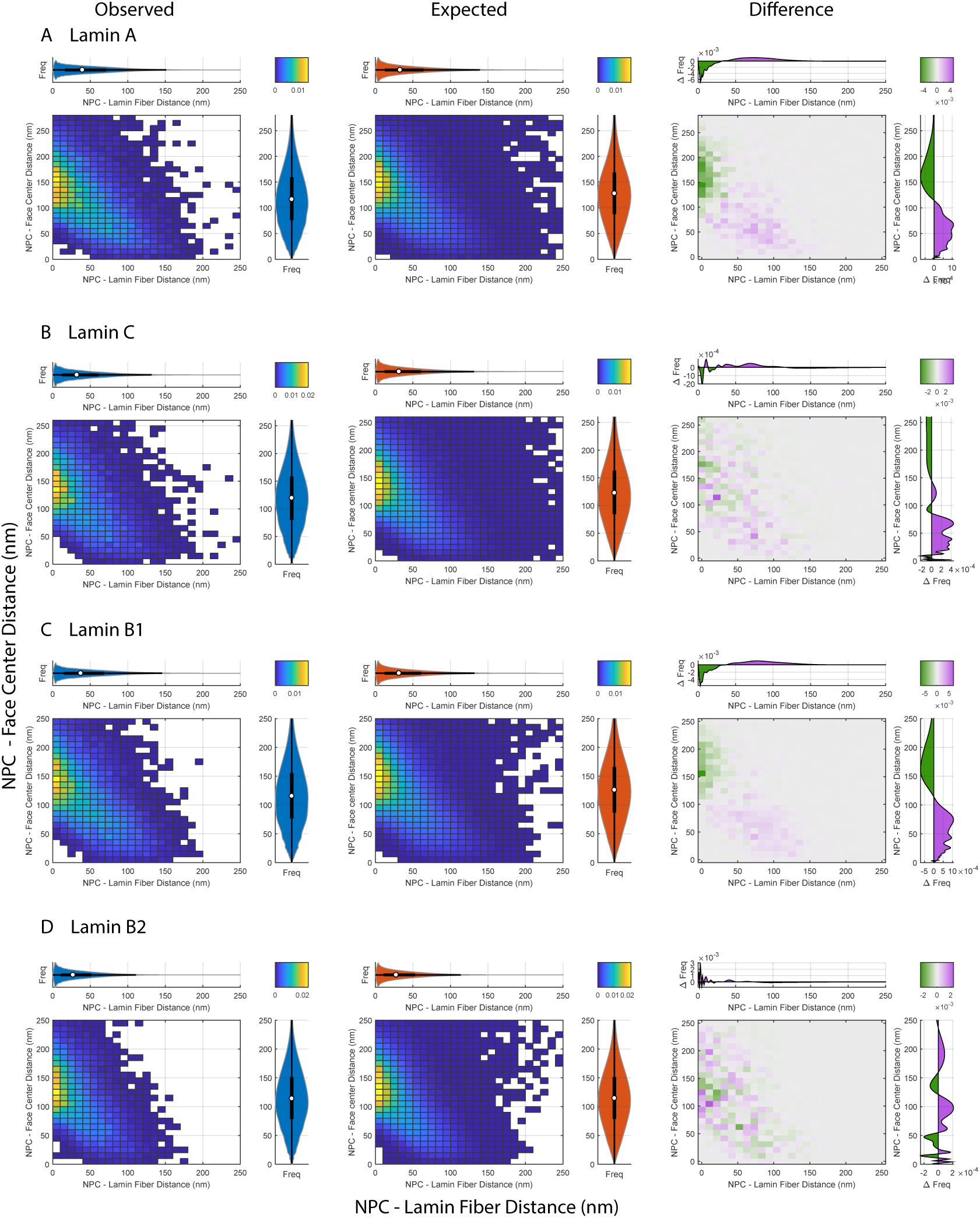
Bivariate histograms of WT MEFs of NPC to face vs fiber distances shows lamin isoform dependent 2D distribution patterns. First column: Observed bivariate distribution, Second column: Expected bivariate distribution under the null hypothesis created by randomizing the positions of NPCs in a Monte Carlo simulation, Third column: Difference between observed and expected bivariate distributions. A) First row shows a bivariate distribution of NPC to Lamin A fiber and face center distances in WT MEFs. B) Second row shows bivariate distributions of NPC to Lamin C fiber and face center distances. C) Third row shows bivariate distributions of NPC to Lamin B1 distances. D) Fourth row shows bivariate distributions of NPC to Lamin B2 distances. First column represents the observed bivariate distribution. Second column represents the expected bivariate distribution. Third column represents the difference between expected and observed. Difference between the observed and expected distance distributions with purple indicating where the observed exceeds the expected frequency and green showing when the observed frequency is less than the expected frequency. Marginal violin plots and box plots of the distances correspond with the half-violin plot counterparts of the same orientation and color as in Panel B of Figure 3.

**Figure S3.**
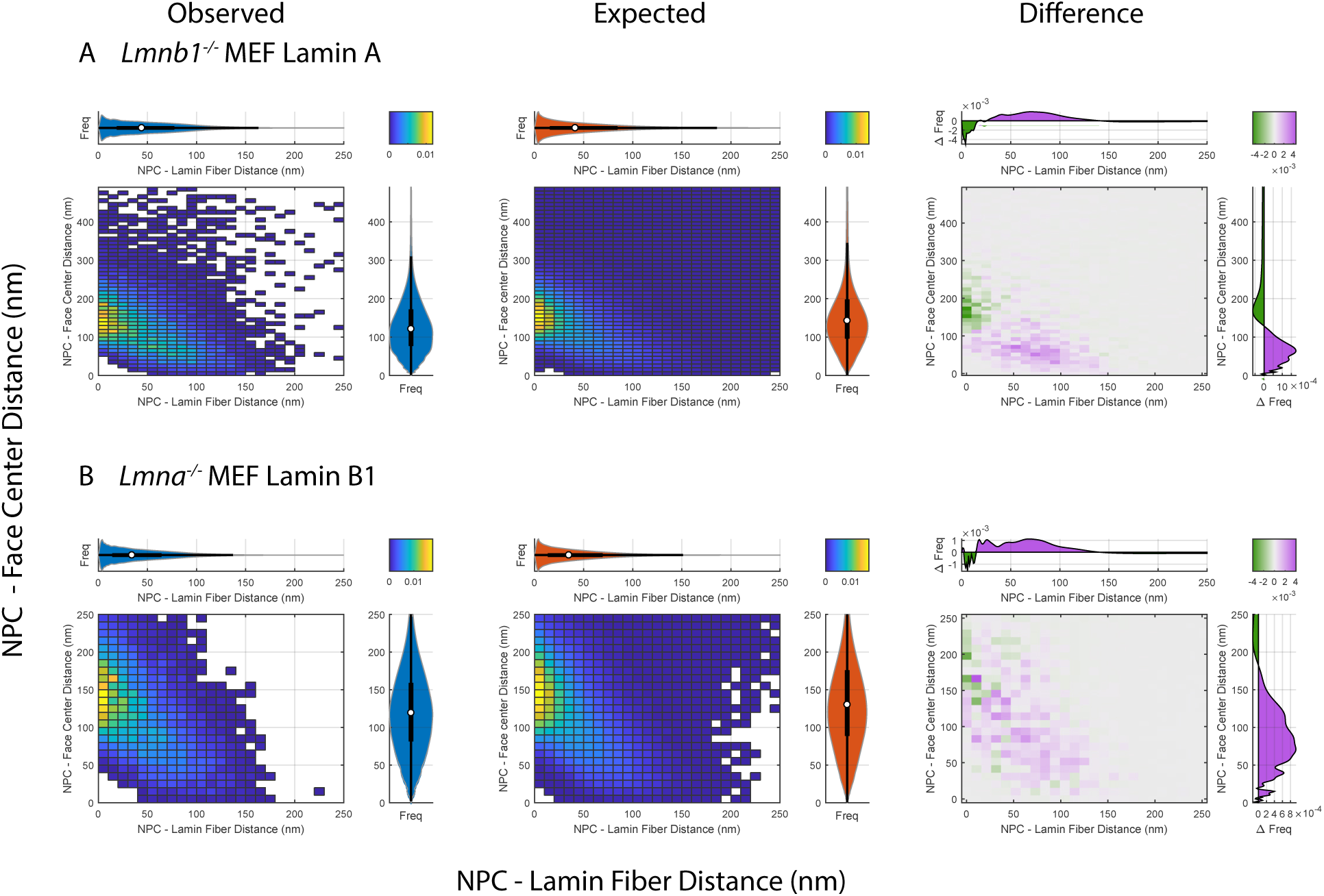
Bivariate histograms of *Lmnb1*^*-/-*^ and *Lmna*^*-/-*^ MEFs. A) First row corresponds NPC to Lamin A fiber and face center distances in *Lmnb1*^*-/-*^ MEFs. B) Second row shows NPC to Lamin B1 fiber and face center distances in *Lmna*^*-/-*^ MEFs. Columns are as in Figure S2.

**Figure S4.**
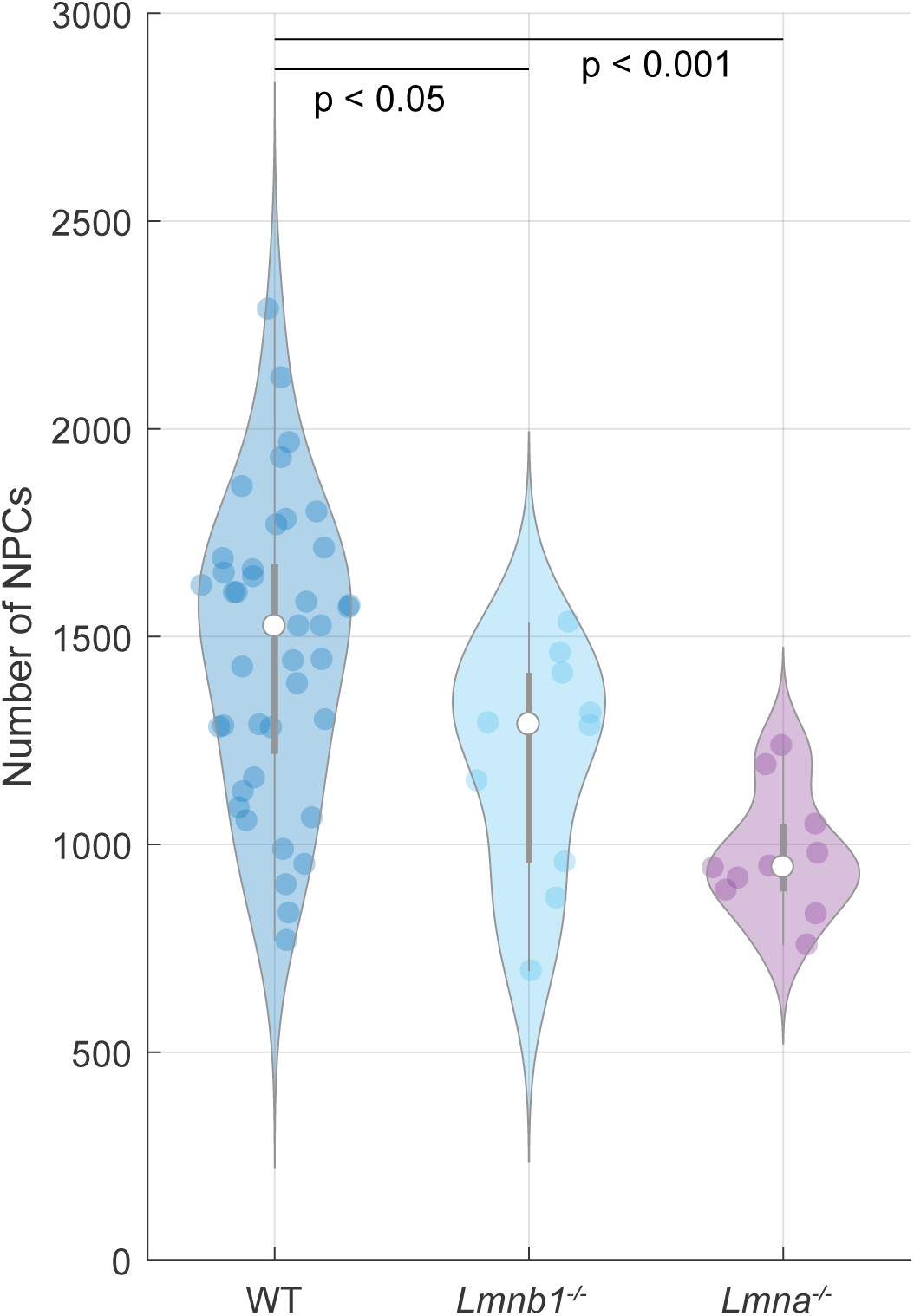
Violin plots comparing the number of NPCs detected in WT, *Lmnb1*^*-/-*^, and *Lmna*^*-/-*^ MEFs. Number of NPCs per cell for WT, *Lmnb1*^*-/-*^, and *Lmna*^*-/-*^ MEFs. The WT category consist of 40 cells pooled from the cells counted in the first four rows of Tables 1A and 1B consisting of cells of WT genotype and stained with antibodies against the four lamin isoforms. The *Lmnb1*^*-/-*^ category consists of 10 cells corresponding to the fifth row of Tables 1A and 1B. The *Lmna*^*-/-*^ category consists of 10 cells corresponding to the sixth row of Tables 1A and 1B. The white circles indicate the medians. The thick grey bar indicates the interquartile range (IQR). The grey whiskers indicate 1.5 times the IQR. Each colored circle corresponds to a single cell. The Mann-Whitney U test was used to compare distributions and determine p-values as described in the Materials and Methods.

**Figure S5.**
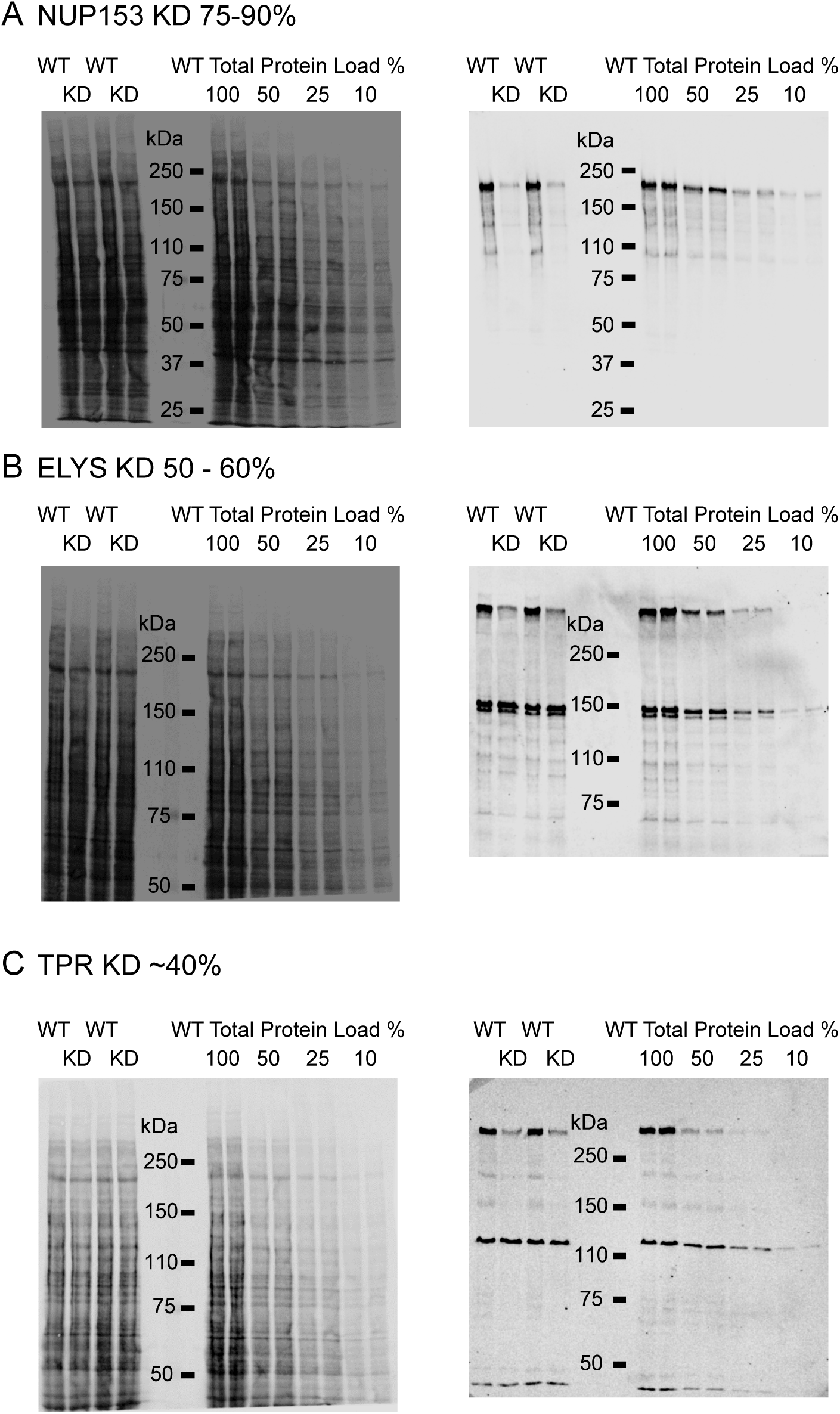
Western blots of ELYS, NUP153, and TPR siRNA knockdown experiments. siRNA knockdowns were carried out and quantified as described in Materials and Methods. The panels on the left are the total protein stains of the immunoblots with each sample loaded in duplicate. The panels on the right are the immunoblots for each antibody A) NUP153, B) ELYS, C) TPR. The degree of knockdown for each protein was determined by quantifying the average intensity of each duplicate after correction for protein load and comparison to the dilution series of the total protein load from WT cells.

**Figure S6.**
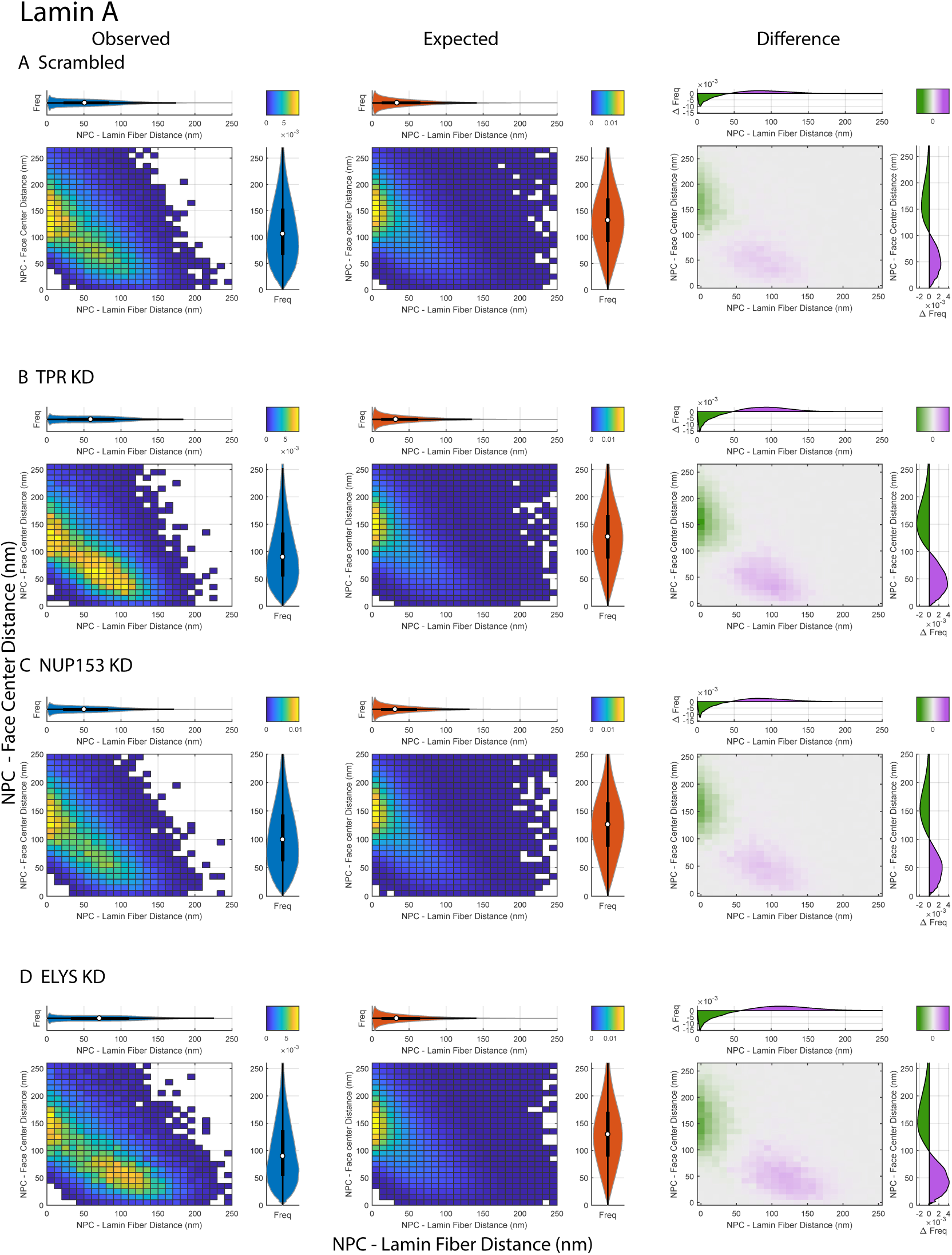
Bivariate histograms of LA fiber-NPC and face center-NPC distances. First column: Observed bivariate distribution, Second column: Expected bivariate distribution under the null hypothesis created by randomizing the positions of NPCs in a Monte Carlo simulation, Third column: Difference between observed and expected bivariate distributions. A) First row shows a bivariate distribution of NPC to Lamin A fiber and face center distances in WT MEFs after treatment with scramble siRNA. B) Second row shows the same with siRNA knockdown of TPR. C) Third row shows the same with siRNA knockdown of Nup153. D) Fourth row shows the same with siRNA knockdown of Elys. First column represents the observed bivariate distribution. Second column represents the expected bivariate distribution. Third column represents the difference between expected and observed. Difference between the observed and expected distance distributions with purple indicating where the observed exceeds the expected frequency and green showing when the observed frequency is less than the expected frequency. Marginal violin plots and box plots of the distances correspond with the half-violin plot counterparts of the same orientation and color as in Panels B-E of Figure 5.

**Figure S7.**
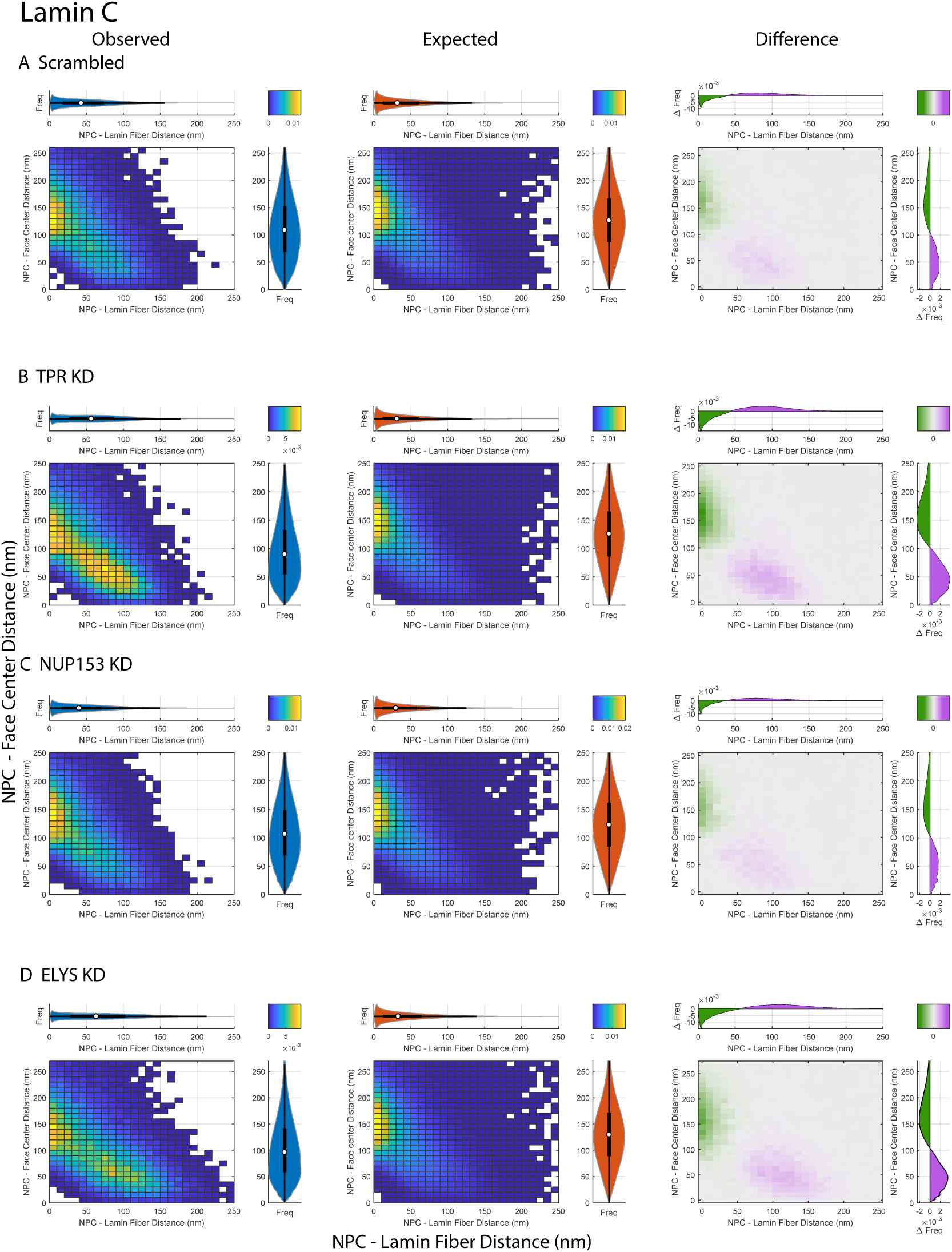
Bivariate histograms of LC Fiber-NPC and Face Center-NPC Distances. First column: Observed bivariate distribution, Second column: Expected bivariate distribution under the null hypothesis created by randomizing the positions of NPCs in a Monte Carlo simulation, Third column: Difference between observed and expected bivariate distributions. A) First row shows a bivariate distribution of NPC to Lamin C fiber and face center distances in WT MEFs after treatment with scramble siRNA. B) Second row shows the same with siRNA knockdown of TPR. C) Third row shows the same with siRNA knockdown of Nup153. D) Fourth row shows the same with siRNA knockdown of Elys. First column represents the observed bivariate distribution. Second column represents the expected bivariate distribution. Third column represents the difference between expected and observed. Difference between the observed and expected distance distributions with purple indicating where the observed exceeds the expected frequency and green showing when the observed frequency is less than the expected frequency. Marginal violin plots and box plots of the distances correspond with the half-violin plot counterparts of the same orientation and color as in Panels B-E of Figure 6.

**Figure S8.**
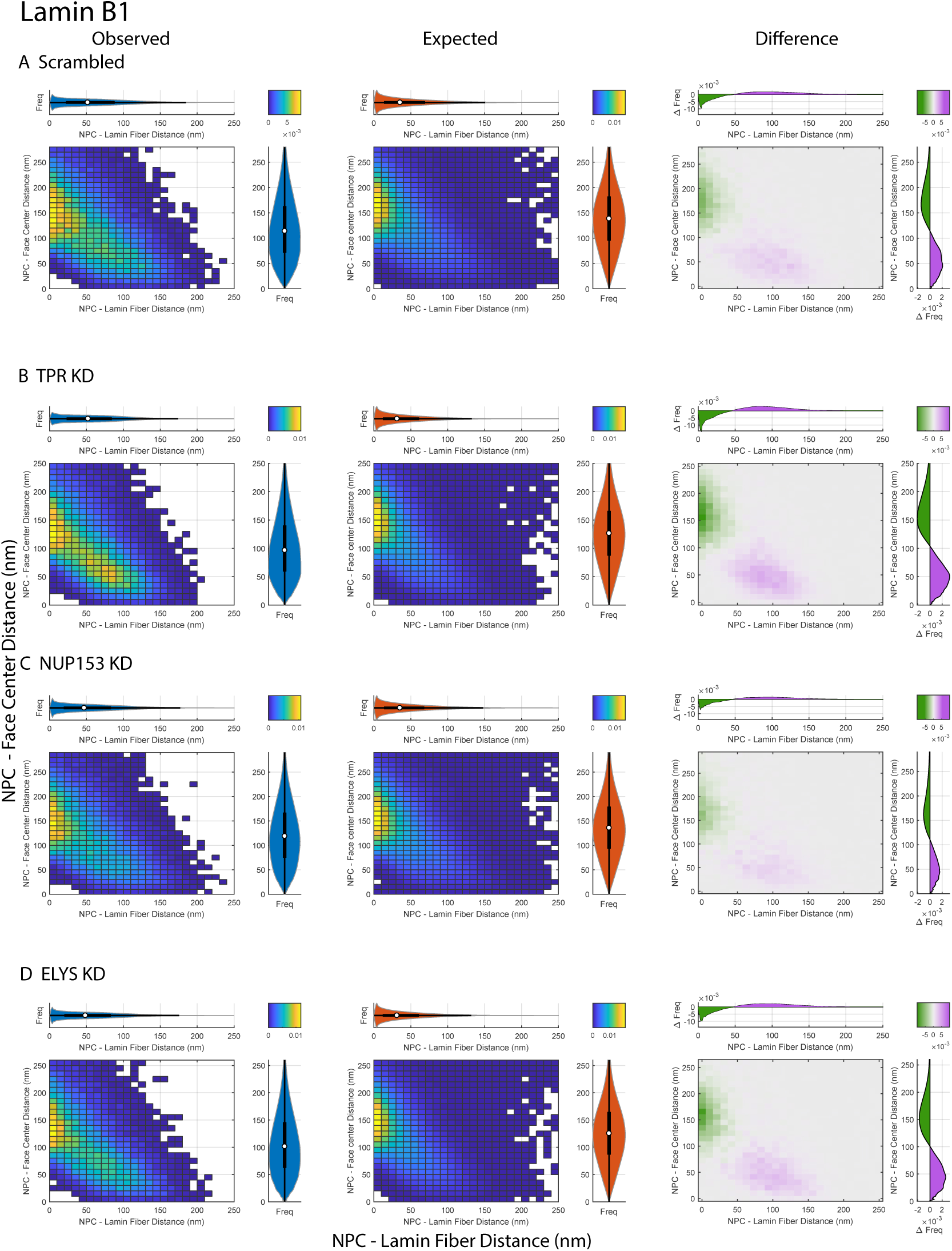
Bivariate histograms of LB1 Fiber-NPC and Face Center-NPC Distances. First column: Observed bivariate distribution, Second column: Expected bivariate distribution under the null hypothesis created by randomizing the positions of NPCs in a Monte Carlo simulation, Third column: Difference between observed and expected bivariate distributions. A) First row shows a bivariate distribution of NPC to Lamin B1 fiber and face center distances inWT MEFs after treatment with scramble siRNA. B) Second row shows the same with siRNA knockdown of TPR. C) Third row shows the same with siRNA knockdown of Nup153. D) Fourth row shows the same with siRNA knockdown of Elys. First column represents the observed bivariate distribution. Second column represents the expected bivariate distribution. Third column represents the difference between expected and observed. Difference between the observed and expected distance distributions with purple indicating where the observed exceeds the expected frequency and green showing when the observed frequency is less than the expected frequency. Marginal violin plots and box plots of the distances correspond with the half-violin plot counterparts of the same orientation and color as in Panels B-E of Figure 7.

**Figure S9.**
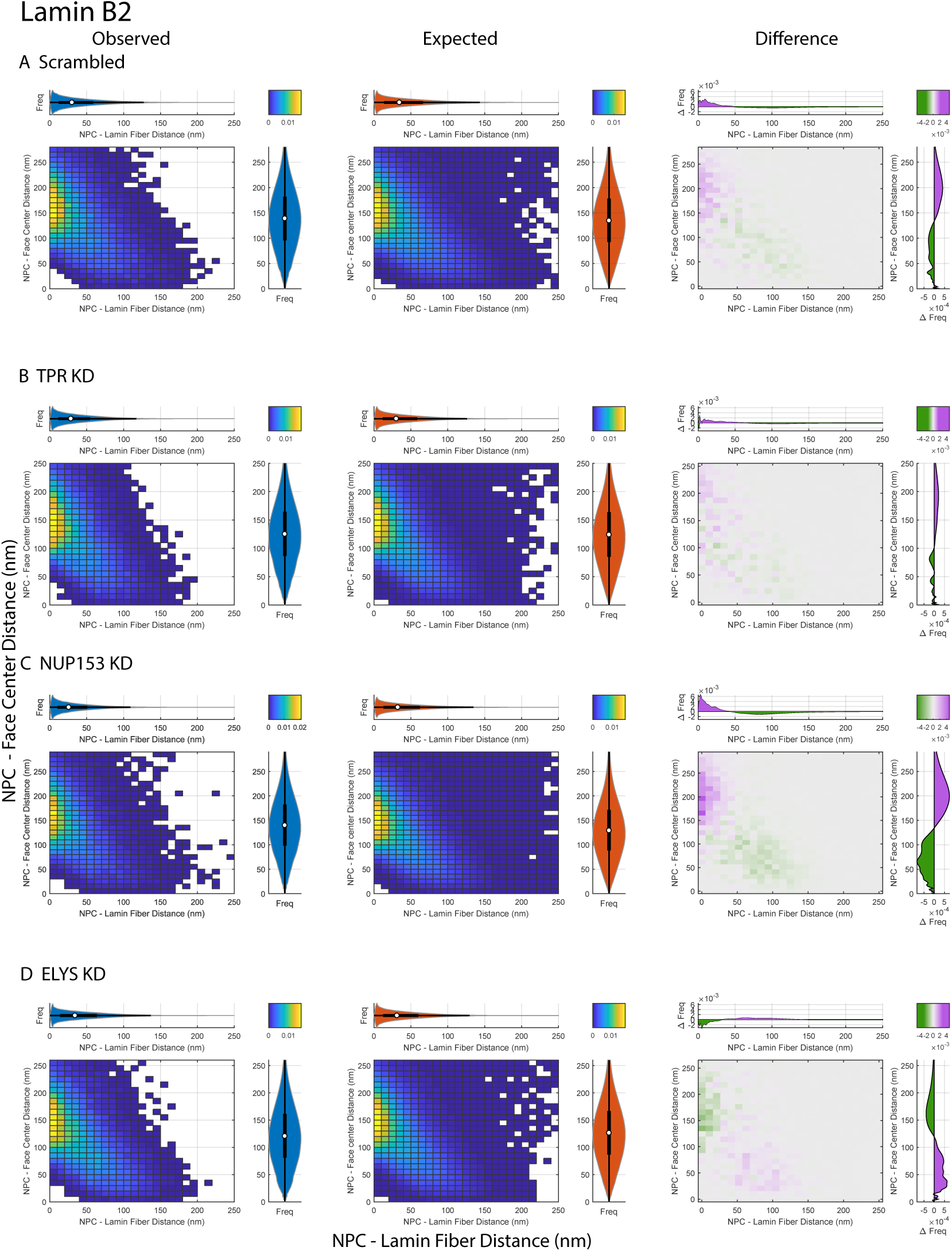
Bivariate histograms of LB2 Fiber-NPC and Face Center-NPC Distances. First column: Observed bivariate distribution, Second column: Expected bivariate distribution under the null hypothesis created by randomizing the positions of NPCs in a Monte Carlo simulation, Third column: Difference between observed and expected bivariate distributions. A) First row shows a bivariate distribution of NPC to Lamin B2 fiber and face center distances in WT MEFs after treatment with scramble siRNA. B) Second row shows the same with siRNA knockdown of TPR. C) Third row shows the same with siRNA knockdown of Nup153. D) Fourth row shows the same with siRNA knockdown of Elys. First column represents the observed bivariate distribution. Second column represents the expected bivariate distribution. Third column represents the difference between expected and observed. Difference between the observed and expected distance distributions with purple indicating where the observed exceeds the expected frequency and green showing when the observed frequency is less than the expected frequency. Marginal violin plots and box plots of the distances correspond with the half-violin plot counterparts of the same orientation and color as in Panels B-E of Figure 8.

**Figure S10.**
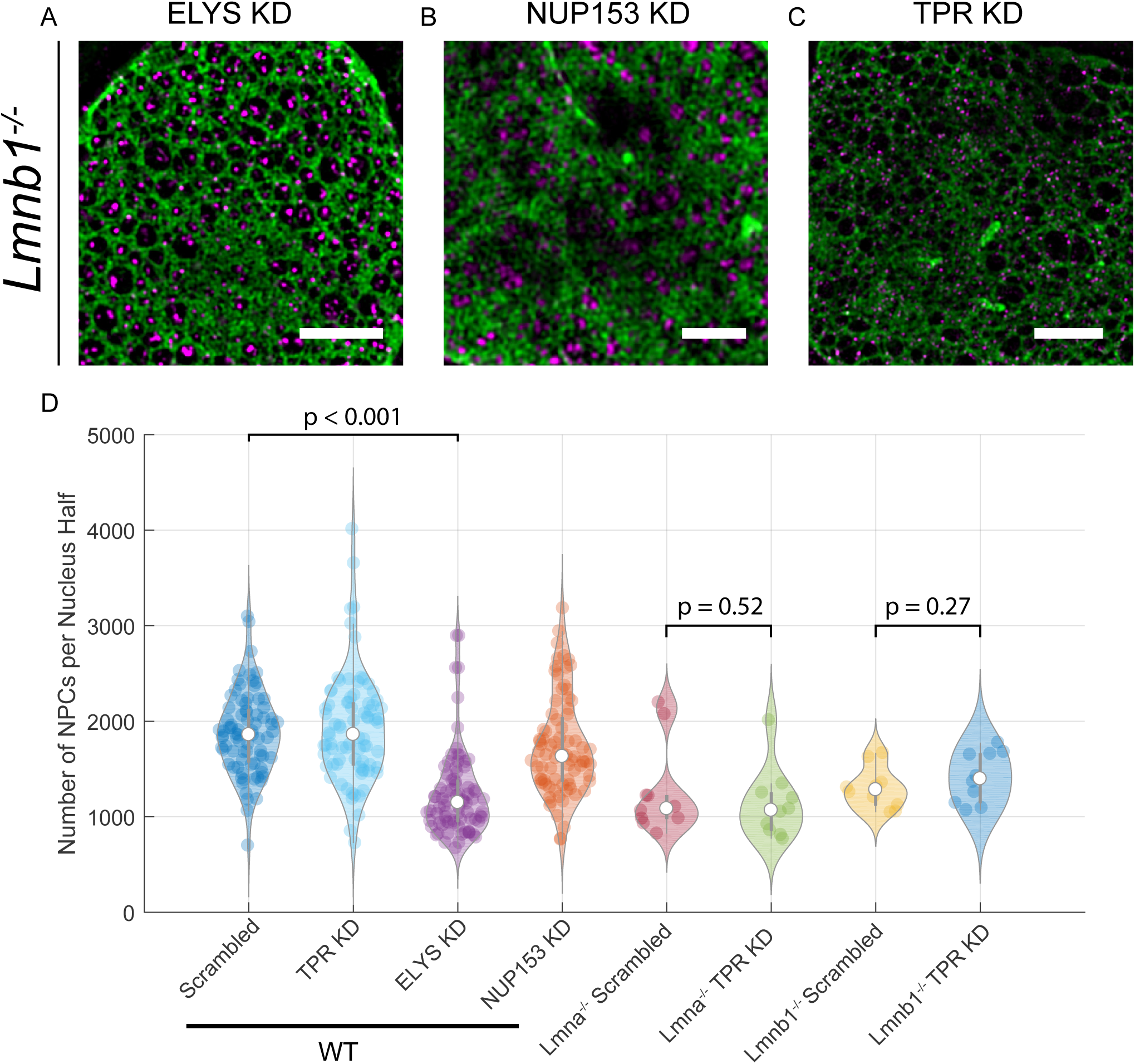
Effect of ELYS, NUP153, and TPR KD in *Lmnb1*^*-/-*^ and *Lmna*^*-/-*^ MEFs. A-C) *Lmnb1*^*-/-*^ MEFs A) ELYS knockdown, B) NUP153 knockdown, C) TPR knockdown, D) Number of NPCs per MEF nuclei in a single focal plane in WT MEFs after ELYS (80 cells), NUP153 (80 cells), and TPR knockdown (80 cells); in *Lmna*^*-/-*^ MEFs after TPR knockdown (10 cells); and in *Lmnb1*^*-/-*^ MEFs after TPR knockdown (10 cells) in comparison to scrambled siRNA (80 WT MEFs, 10 *Lmna*^*-/-*^ MEFs, 10 *Lmnb1*^*-/-*^ MEFs). The white circles indicate the medians. The thick grey bar indicates the interquartile range (IQR). The grey whiskers indicate 1.5 times the IQR. Each colored circle represents a single cell. The Mann-Whitney U test was used to compare the distributions as described in the Materials and Methods. Scale Bar = 10 *µm*

